# Transcript-Specific Site-Directed RNA Editing Reveals Principles for Splice-aware guide RNA Design

**DOI:** 10.64898/2026.07.30.741163

**Authors:** Nina Schneider, Nave Zehoray, Ricky Steinberg, Eyal Banin, Yvan Arsenijevic, Shay Ben Aroya, Erez Y. Levanon, Dror Sharon

**Affiliations:** Division of Ophthalmology, Hadassah Medical Center, Faculty of Medicine, The Hebrew University of Jerusalem, Jerusalem, Israel; The Mina and Everard Goodman Faculty of Life Sciences, Bar-Ilan University, Ramat Gan, Israel; Group of Retinal Degeneration and Regeneration, Department of Biomedical Sciences, University of Lausanne, Lausanne, Switzerland

**Keywords:** RNA editing, ADAR, Splicing, guideRNA design, Exonic Splice Enhancers

## Abstract

Site-directed RNA editing (SDRE) utilizing the adenosine deaminase acting on RNA (ADAR) enzymes is commonly facilitated by guide RNAs (gRNAs) optimized to enhance on-target editing and minimize bystander effects. However, the impact of gRNA binding and ADAR-mediated SDRE on canonical pre-mRNA splicing remains poorly understood. Here, we developed an in vitro transcript-specific editing strategy that enables selective targeting and direct comparison of SDRE in pre-mRNA and mature mRNA. Using splice-relevant variants associated with inherited retinal diseases, we investigated the effects of SDRE on exonic, near-canonical intronic, and deep intronic splice variants. We identified gRNA-induced splice perturbation at exonic and intronic targets and observed that higher editing levels could be associated with increased splice disruption. Conversely, SDRE of two exonic splice variants and a deep intronic variant resulted in increased production of correctly spliced transcripts, demonstrating the potential of SDRE for splice modulation. Finally, by dissecting the effects of ADAR expression and gRNA design on editing and splicing outcomes, we established a system for identifying design principles that reduce splice interference and enhance the generation of correctly spliced, edited transcripts. These findings highlight the importance of considering transcript context and splicing consequences in the development of SDRE-based therapeutic strategies.

**GRAPHICAL ABSTRACT:** Created in BioRender. Schneider, N. (2026) https://BioRender.com/59i0tiy

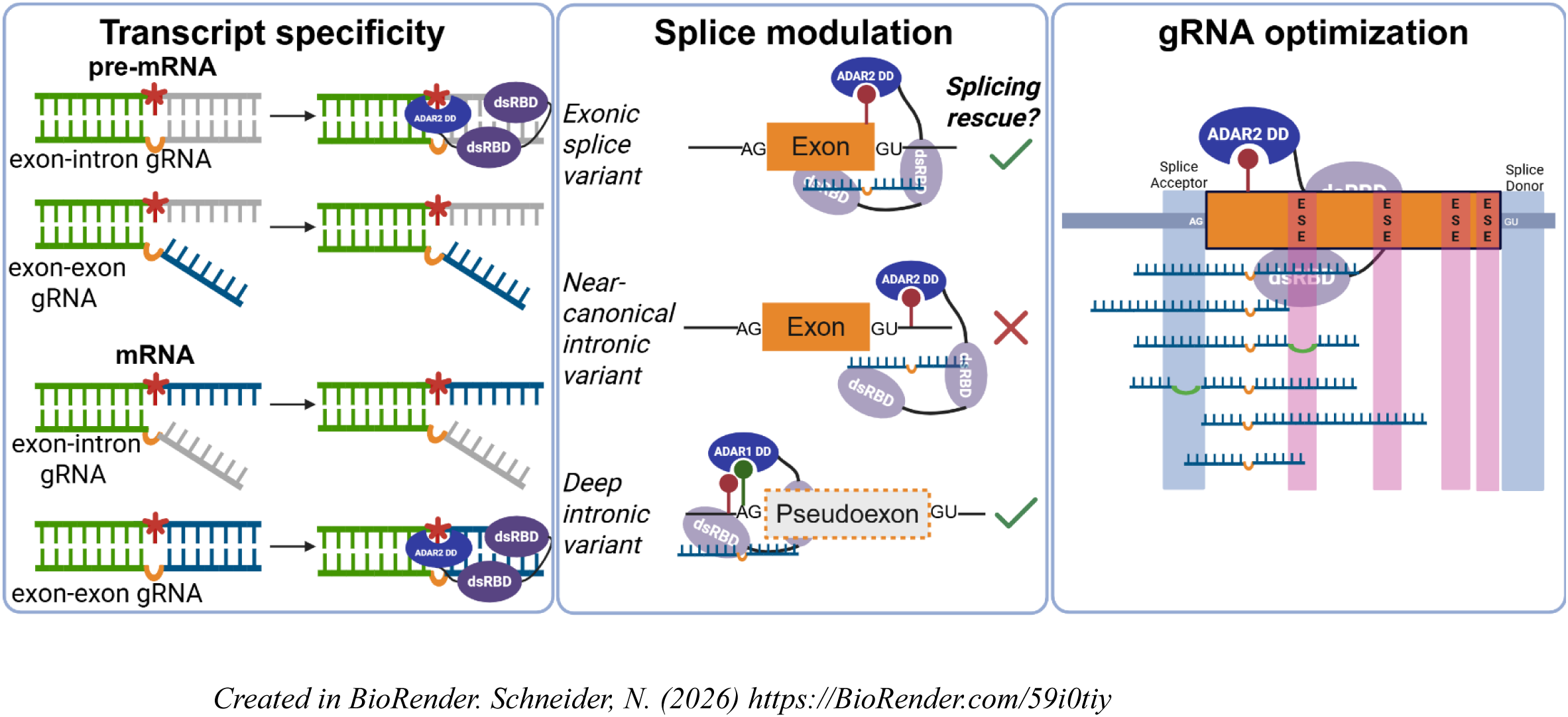

## INTRODUCTION

Site-directed RNA editing (SDRE) using adenosine deaminases acting on RNA (ADARs) is a transient and sequence-specific approach for correcting pathogenic variants at the RNA level through adenosine-to-inosine (A-to-I) editing. In mammalian cells, inosine is generally read as guanosine during translation, so A-to-I RNA editing can be treated functionally as an A-to-G substitution (1). Cellular RNA editing is mediated by three ADAR isoforms: the predominantly nuclear ADAR1 p110, the interferon-inducible ADAR1 p150 which shuttles between the nucleus and cytoplasm, and the predominantly nuclear ADAR2 (2–5). Numerous strategies have been developed to efficiently recruit the ADAR1 p110 and ADAR2 enzymes to a target adenosine, generally in exonic regions, using exogenous guide RNAs (gRNAs) to artificially create double-stranded RNA (dsRNA). These approaches have explored diverse gRNA designs, including variations in guide length (6), symmetry (7), recruitment sequences (8, 9), mismatches (10), bulges (11–13), chemical modifications (14), and structure (15, 16) with the primary goals of maximizing on-target editing while minimizing bystander editing. However, comparatively little attention has been paid to the transcript context in which SDRE occurs, the consequences of gRNA-mediated editing for pre-mRNA splicing, or the application of SDRE to splice-related variants in exonic and intronic regions as a means of splice modulation.

Endogenous editing of the transcriptome by the ADAR enzyme has been shown to take place co-transcriptionally on nascent RNA before polyadenylation (17). The timing and choreography of ADAR-mediated editing play a critical role in gRNA design; targeting the pre-mRNA or mRNA depending on the adenosine target location’s proximity to the exon-intron boundary and length of the gRNA. Few have addressed this interplay in relation to gRNA design by comparing SDRE levels between pre-mRNA and mRNA (6, 12), and oftentimes the gRNAs differ only within a minor region of the sequence or mRNA editing levels are measured as what remains after the pre-mRNA has been edited, leaving it unclear whether pre-mRNA and mature mRNA can be selectively and independently targeted by SDRE.

Pre-mRNA splicing has been shown to be influenced by endogenous ADAR activity through multiple mechanisms. Studies suggest that A-to-I editing may modulate splicing by creating novel 3’ splice sites (17–19), modifying cis-acting elements (20, 21) and disrupting branch point sequences (22) as well as changing RNA secondary structure (23, 24). Splice sites have also been noted to be disrupted by RNA editing, for example, weakening the predicted strength of a 3’ splice site (17). Though there is extensive evidence that ADAR can splice-modulate through its RNA editing activity, few studies have utilized this evidence for targeted editing for the purpose of splice modulation (25, 26).

In addition to the editing-dependent effect RNA editing by ADAR has on splicing patterns, studies have shown a possible editing-independent role that ADAR has in splicing modulation as well (27–31). One study showed through mutant HEK293T and EC109 cell lines devoid of either the ADAR enzymatic activity or dsRNA-binding capabilities that recruiting ADAR to an exon-intron junction blocks exon recognition by the spliceosome, while binding of ADAR to intra-intronic dsRNA may promote exon inclusion by enhancing intron definition (29). These observations suggest that recruitment of ADAR by exogenous gRNAs during SDRE may influence splicing independently of RNA editing.

Splice-related effects of gRNA binding during SDRE may also arise independently of ADAR activity and instead reflect the intrinsic characteristics of exons. Most internal human exons are relatively short (<300 nucleotides; average ∼150 nucleotides) (32) and are densely populated with cis-acting regulatory elements, including exonic splice enhancer (ESE) motifs that contribute to both constitutive and alternative splicing through binding of SR proteins (33, 34). Accordingly, even a 60-nt gRNA targeting an exon is likely to overlap multiple putative SR protein-binding sites (Figure 1A). In addition, splice donor and acceptor sites require coordinated recognition by the spliceosome and associated regulatory factors and may be sterically disrupted by gRNA binding, potentially leading to exon skipping (35, 36) (Figure 1A). Together, these features suggest that gRNAs designed primarily for efficient SDRE may inadvertently perturb splicing, highlighting ESE distribution and splice site locations as critical yet underappreciated determinants of SDRE design.

**Figure 1.**
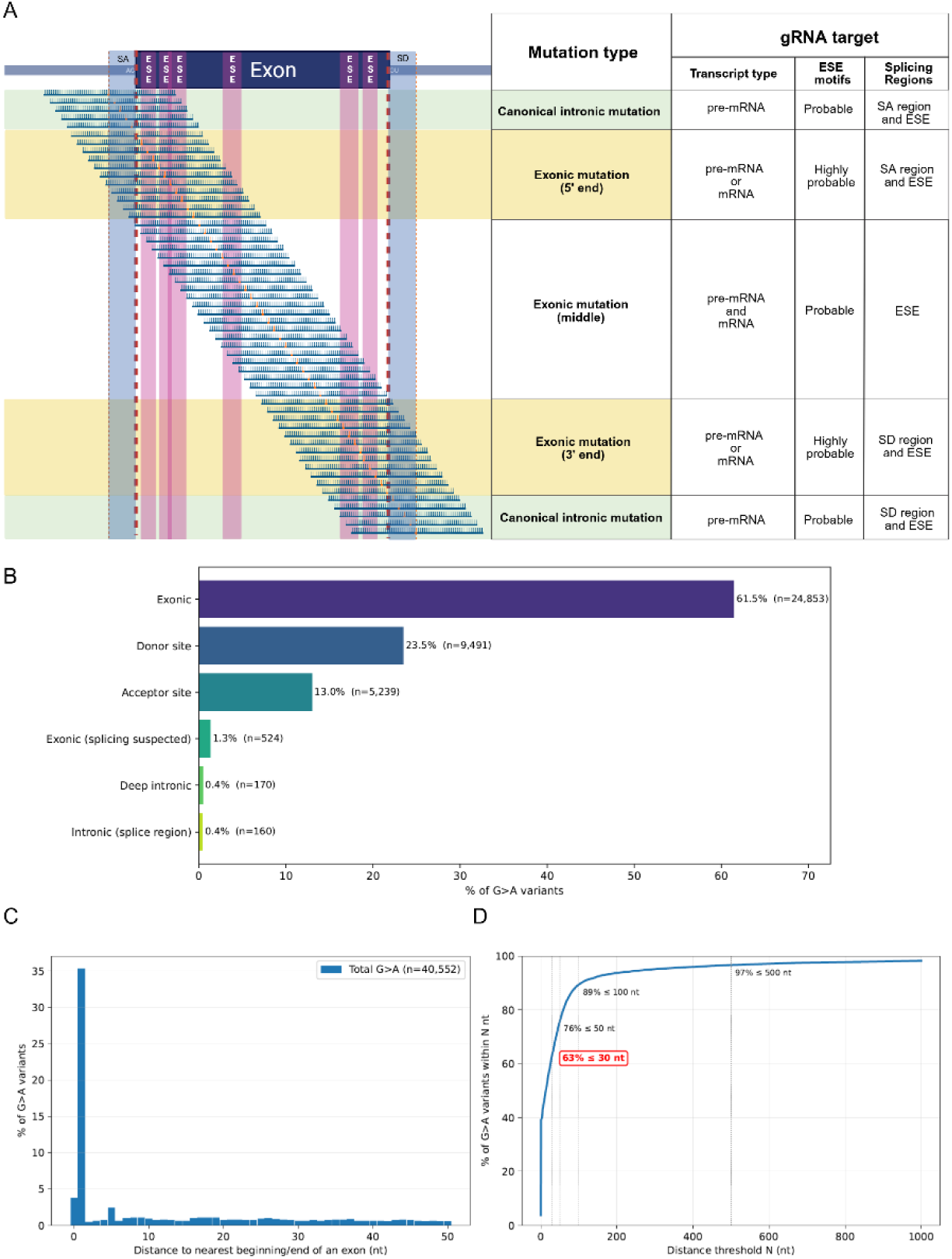
Target position determines transcript context and splice-related features encountered by SDRE gRNAs. (A) Schematic representation of representative 60-nt symmetric gRNAs targeting different regions of a transcript, including the middle of an exon, exon termini, and intronic regions. Depending on target location, gRNAs may overlap exonic splicing enhancer (ESE) motifs, splice donor (SD) or splice acceptor (SA) sites, and may require consideration of pre-mRNA versus mature mRNA targeting. Orange lines indicate the orphan cytosine positioned opposite the target adenosine for ADAR-mediated A-to-I editing. SA, splice acceptor; SD, splice donor; ESE, exonic splicing enhancer motifs. (B-D) Pathogenic (P) and likely-pathogenic (LP) G>A variants in ClinVar (release 20260715) within a MANE Select transcript, exonic and intronic (n = 40,552); G>A is defined on the sense strand (See Methods). (B) Variants grouped into splice classes: exonic-not predicted to affect splicing; donor and acceptor sites-intronic variants within 5 nt of the 5′ or 3′ splice site, respectively; exonic (splicing suspected)-exonic variants with a SpliceAI delta score ≥ 0.8; deep intronic-more than 30 nt into the intron; intronic (splice region)-6-30 nt into the intron from either junction. Intronic classes are defined by position alone. Exonic variants without a SpliceAI score could not be assigned an exonic class and are excluded from this panel. (C, D) Distance of each variant from the nearest canonical splice junction (defined as the first or last base of an exon). (C) Percentage of variants at each exact distance, 0-50 nt. (D) Cumulative percentage of variants within N nt of the nearest junction, 0-1,000 nt; dotted lines mark 30, 50, 100 and 500 nt, and the value at 30 nt is boxed in red. *Created in BioRender. Schneider, N.* (*2026*) https://BioRender.com/fzfs3et

In this study, using inherited retinal disease-associated variants as targets, we investigated the relationship between transcript type, ADAR-mediated SDRE, and pre-mRNA splicing. We developed a transcript-specific editing strategy to directly compare editing of pre-mRNA and mature mRNA and determine whether each transcript type can be selectively targeted by exogenous gRNAs. Once we established that one can exclusively target pre-mRNA, we studied the effects of SDRE on exonic, intronic, and deep intronic splice-affecting variants. Finally, by examining the effects of ADAR expression and gRNA binding on RNA editing and splicing at exonic targets, we established gRNA design principles that minimize splice disruption while maximizing the production of correctly spliced, edited transcripts.

## MATERIALS AND METHODS

### Bioinformatic Analysis

The methods presented in this section refer to Figure 1 B-D.

Variant set-Pathogenic and likely-pathogenic (P/LP) single-nucleotide variants were obtained from the dated ClinVar release `clinvar_20260715` (GRCh38) (37). Records were retained when `CLNSIG` contained the exact token `Pathogenic` or `Likely_pathogenic`, matched after splitting on `/`,, and `|`, so that `Conflicting_classifications_of_pathogenicity` and `Variants_of_uncertain_significance` were excluded. Alternate alleles were processed individually, and variants were identified by (chromosome, position, reference, alternate). Analysis was restricted to the canonical nuclear chromosomes (chr1-22, X, Y); 118 mitochondrial variants were excluded, as mitochondrial transcripts are not spliced by the spliceosome. This yielded 189,109 P/LP single-nucleotide variants.

Transcript scope-All analyses use MANE Select transcripts (38) from GENCODE v50, Ensembl 116 (39). Each variant was assigned a region - exonic, intronic, or intergenic / outside MANE - by intersection with MANE Select exons and transcript spans. The analysis cohort comprises the exonic and intronic variants (n = 188,621); 468 variants (0.25%) fell outside any MANE Select transcript and were excluded, as were the strand-ambiguous variants defined below.

Definition of a G>A variant-Because ADAR acts on the transcribed strand, G>A was defined on the sense strand: a genomic G>A in a plus-strand gene, or a genomic C>T in a minus-strand gene. Strand was assigned by a strict cascade: (i) the MANE Select transcript containing the variant; (ii) where a variant lay within two antisense-overlapping MANE transcripts, or outside MANE, the curated gene or genes reported for that variant in ClinVar `variant_summary_2026-07` were each mapped to their GENCODE strand (HGNC identifier preferred over gene symbol). The strand was accepted only when every listed gene indicated the same strand. (iii) Conflicting or absent curation left the strand unresolved (54 variants). Unresolved variants whose substitution is a genomic G>A or C>T - the only substitutions whose sense-strand identity depends on strand - could not be classified and were excluded from all totals (n = 20); the remaining unresolved variants cannot be a sense-strand G>A under either strand and were retained as non-G>A. This yielded 40,552 sense-strand G>A variants in the cohort.

Splice-site geometry. Canonical splice junctions were derived from MANE Select exon boundaries: internal exons contribute two junctions, first and last exons contribute only their intron-facing edge, and single-exon transcripts contribute none. Junctions were deduplicated to unique genomic positions and labelled donor or acceptor in a strand-aware manner, giving 365,637 junctions. The junction was anchored at the exon’s terminal base, so that an intronic variant at HGVS position `c.X+5` lies 5 nt from the junction. Distances are genome-level signed distances from each variant to the nearest junction, computed with `bedtools closest -D ref` (40); magnitudes are reported.

Splice-effect prediction-Predicted splicing effects were taken from the precomputed Ensembl SpliceAI MANE v1.4 delta scores (SpliceAI v1.3.1) (41), queried by chromosome, position and alternate allele with tabix (42). For each variant we recorded the four delta scores (acceptor gain, acceptor loss, donor gain, donor loss). Where a variant’s position and allele carried entries for more than one gene, the entry annotated with the variant’s own gene was used - the MANE Select gene containing it on the sense strand assigned above. A variant’s prediction and its G>A status therefore derive from the same gene. Where the variant’s gene was unresolved, or none of the entries named it, no prediction was assigned rather than substituting another gene’s value. Variant positions carrying a single entry were used as given. The per-variant effect is the maximum of the four deltas, and a delta ≥ 0.8 was treated as a strong predicted effect throughout.

Splice classes-Each variant was assigned to exactly one class: *Donor site* and *Acceptor site* (intronic, within 5 nt of the respective junction), *Intronic (splice region)* (6-30 nt into the intron from either junction), *Deep intronic* (more than 30 nt into the intron from either junction), *Exonic (splicing suspected)* (exonic, per-variant delta ≥ 0.8), and *Exonic*. A variant was counted as splice-associated if it fell in any class other than *Exonic*.

Missing predictions-The Ensembl SpliceAI MANE precompute does not carry an entry for every MANE transcript: 985 cohort variants (726 exonic, 259 intronic) have no score, concentrated in a subset of MANE genes and in non-coding MANE genes. Because only the exonic classes are defined by a score, the 115 unscored exonic G>A variants could not be classified and were excluded from the splice-class breakdown (n = 40,437 of 40,552) rather than assigned to a class; positional classes and all distance analyses retain every variant. For quantities whose definition requires a prediction, the accompanying Supplementary Table 1 gives bounds in which the lower bound treats every unscored variant as unaffected and the upper bound treats all of them as affected: the splice-associated fraction is 38.5% [38.5-38.7%] and the *Exonic (splicing suspected)* class 1.3% [1.3-1.6%]. Quantities defined by position are exact and carry no bounds.

Software and reproducibility-Analyses were run in the pinned conda environment supplied with the code (Python 3.12, pandas 2.3.3, numpy 2.0.2, matplotlib 3.9.4, bedtools 2.31.1, HTSlib 1.19), which the build places on PATH so a run cannot pick up a different toolchain. Input versions are pinned by dated filename and verified against provider checksums where published, with provenance recorded in a tracked manifest. The complete pipeline runs from raw downloads to figures as a single command, every analysis parameter resides in one configuration file, and each parser carries self-tests.

### Guide RNA design

Chemically modified gRNAs ranging from 22 to 60 nucleotides in length were synthesized by Integrated DNA Technologies (IDT). All gRNAs contained three 2′-O-methyl-modified nucleotides and three phosphorothioate linkages at both the 5′ and 3′ termini. A cytosine mismatch was incorporated opposite the target adenosine (or “orphan” base) in all designs.

Depending on sequence-specific synthesis constraints, gRNAs were obtained either as HPLC-purified or standard desalted oligonucleotides. Individual gRNA sequences, lengths, purification method, and synthesis scale are provided in Supplementary Table 2.

Putative ESE motifs were identified using ESEfinder 3.0 (43, 44) with default parameters. Predicted SR protein-binding sites (SRSF1, SRSF2, SRSF5, and SRSF6) were mapped in regions surrounding target variants and used to influence gRNA design to avoid disruption of predicted splice-regulatory elements for some gRNAs as described in the text.

### Plasmid design

Reporter plasmids for minimal or no-splicing assays for *ABCA4* c.4634G>A and *ABCA4* c.5196+1137G>T were generated by restriction-ligation cloning of synthetic gBlocks (Integrated DNA Technologies; 163-166 nt prior to restriction). Inserts contained a 60-nt *ABCA4*-derived sequence harboring the respective variant and were cloned in-frame into a dual fluorescent reporter plasmid expressing mCherry and EGFP (Addgene #86639, Watertown, MA, USA). Restriction digestion was performed using KpnI (Thermo Fisher Scientific #00914286) and BamHI-HF (New England Biolabs #R3136), followed by ligation and transformation into DH5α-HIT chemically competent cells (RBC #RH618, New Taipei City, Taiwan). Plasmids were purified using a QIAprep Spin Miniprep Kit (Qiagen #2714) and verified by Sanger sequencing. gBlock sequences are listed in Supplementary Table 3.

Minigene splicing reporters for *ABCA4* c.1556G>A, *CDH23* c.5712G>A, *BBS1* c.479G>A, and a wild-type *PDE6B* c.1832A construct were generated by restriction-ligation cloning of larger synthetic gBlocks (700-1000 bp) into the Exontrap pET01 vector (MoBiTec GmbH, Germany) (Supplementary Table 3).

Previously described *ABCA4* midigene constructs corresponding to c.4634G>A, c.5196+1137G>A/T, and c.5714+5G>A were kindly provided by Prof. Frans Cremers and Prof. Rob Collin (referred to as BA21, BA32, and BA27); see (45, 46). The *ABCA4* c.5196+1137G>T construct is identical to BA32 c.5196+1137G>A except for the single G>T substitution in the same location.

To simulate complete A-to-G editing affecting splicing for *ABCA4* c.5196+1137G>T, modified Exontrap constructs were generated by cloning 500-bp gBlocks into pET01 containing either the wild-type sequence, the c.5196+1137G>T variant, or a matched A-to-G-edited mimic (c.5196+1137_1138GA>TG) (Supplementary Table 3).

### Antisense oligonucleotides

A series of overlapping 2′-O-methoxyethyl (2′-MOE) antisense oligonucleotides (17-21 nt; 5 nmol, standard desalting) were designed to tile the last 30 nucleotides of exon 11 and the first 29 nucleotides of exon 12. Oligonucleotides were synthesized by Integrated DNA Technologies (IDT) and used to identify regions in which hybridization disrupted normal splicing (Supplementary Table 4).

### In vitro splicing confirmation

HeLa cells (cultured in high-glucose DMEM supplemented with 10% FBS, 1% L-glutamine, and 1% penicillin-streptomycin) and/or 661W cells (cultured in high-glucose DMEM supplemented with 2.5 mM L-glutamine, 20% FBS, 1% penicillin-streptomycin, 40 μg/L hydrocortisone 21-hemisuccinate, 40 μg/L progesterone, 32 mg/L putrescine, and 0.004% β-mercaptoethanol) (47) were seeded at 4×10⁴ cells per well in a 24-well plate. Twenty-four hours after seeding, media was replaced and splice plasmids (0.5 μg/well) were transfected using TransfeX (ATCC, Manassis, VA, USA; Cat. ACS-4005). Media was replaced again 48 h after seeding, and cells were harvested 72 h after seeding for RNA isolation using the NucleoMag™ RNA Kit (Macherey-Nagel, GmbH, Germany; Cat. No. 744350.4). cDNA was synthesized using the qScript cDNA Synthesis Kit (QuantaBio, Beverly, MA, USA; Cat. No. 95047).

### In vitro editing in HeLa cells

Stable ADAR1 p110- and ADAR2-overexpressing HeLa cells were cultured and transfected as previously described (48). Briefly, 4×10^4^ cells were seeded per well in a 24-well plate, and ADAR-overexpressing cells were induced with 1 ng/mL doxycycline where indicated. Twenty-four hours after seeding, splice or reporter plasmids (0.5 μg/well) were transfected using TransfeX (ATCC, Manassis, VA, USA; Cat. ACS-4005). Forty-eight hours after seeding, chemically modified gRNAs were transfected using Lipofectamine RNAiMAX (25 pmol per gRNA type; for experiments involving multiple gRNAs, 25 pmol of each gRNA type was transfected per well). Cells were harvested 72 h after seeding for RNA isolation using the NucleoMag™ RNA Kit (Macherey-Nagel, GmbH, Germany; Cat. No. 744350.4). cDNA was synthesized using the qScript cDNA Synthesis Kit (QuantaBio, Beverly, MA, USA; Cat. No. 95047). Unlike the previous study, cells were not processed for immunofluorescence, and editing levels were assessed by Sanger sequencing or targeted next-generation sequencing (NGS).

### RNA editing analysis

RNA editing analysis was performed on PCR-amplified cDNA using primers listed in Supplementary Table 5 and Supplementary Table 6. Editing levels were quantified either by Sanger sequencing or NGS, depending on the experiment (Supplementary Table 5). For Sanger sequencing data, A-to-I editing was quantified from chromatogram peak heights at the target site. Editing efficiency was calculated as the ratio of guanine signal to the sum of guanine and adenine signals (G / [A + G]) using ImageJ. For NGS-based analysis, FASTQ files were processed using FastQGroomer and aligned using HISAT2. Editing levels were calculated as the fraction of guanine calls at the target position relative to total adenine and guanine calls. In both Sanger sequencing and NGS, background signal derived from non-edited guanine in control samples was subtracted from final editing values.

For splice isoform-specific analysis, alignments were performed against the corresponding isoform reference sequences in NGS while Sanger sequencing-based quantification of specific splice products were derived from gel-excised PCR bands that were purified prior to sequencing and sequenced using isoform-specific primers (Supplementary Table 5, Supplementary Table 6). In the case of several isoforms containing intron 5 in for *BBS1* c.479G>A, sequencing was performed using a reverse primer located within intron 5 to capture all intron 5-retained products and to enable accurate quantification of editing levels.

### Splicing analysis

Splicing analysis of PCR products was performed using capillary electrophoresis on the Agilent TapeStation D1000 ScreenTape®, and peak areas were automatically quantified using TapeStation Analysis Software v5.1. The percentage of wild-type-length transcripts was calculated as the integrated area of the full-length peak relative to the sum of all detected PCR product peaks below the PCR size threshold. Values were then normalized to the corresponding untreated control sample to account for maximum percent full-length transcripts.

For the *ABCA4* c.5196+1137G>T midigene assay, isoform-specific quantification was performed using droplet digital PCR (ddPCR). Approximately 200 ng of cDNA was analyzed, using ddPCR Supermix for Probes (Bio-Rad, Hercules, CA, USA) with isoform-specific primers and probes (Integrated DNA Technologies; Supplementary Table 7). Droplets were generated and PCR amplification was performed under the following cycling conditions: enzyme activation at 95°C for 10 min (1 cycle), denaturation at 95°C for 30 s followed by annealing/extension at 56°C for 1 min (40 cycles), and enzyme deactivation at 98°C for 10 min (1 cycle). Fluorescence signals were read using the QX200 Droplet Reader (Bio-Rad), and positive/negative droplet thresholds were set manually for each experiment.

Biological triplicates with technical duplicates were performed, and technical replicates were averaged prior to downstream analysis. The proportion of pseudoexon-containing transcripts was calculated as:

Pseudoexon inclusion (%) = PE inclusion / (PE inclusion + exon 36-37 junction transcripts).

### Statistical analysis and reproducibility

Statistical analyses were performed using GraphPad Prism (version 11.0.2). Data are presented as mean ± standard error of the mean (SEM) unless otherwise indicated. Biological replicates represent independent experiments performed on separate days (n=3 in most cases, n=2 in cases of experimental loss). Technical replicates for ddPCR duplicates were averaged prior to statistical analysis. For comparisons between two groups, statistical significance was assessed using Welch’s unpaired two-tailed Student’s t-test. For comparisons involving more than two groups within the same experiment, one-way analysis of variance (ANOVA) followed by Tukey’s multiple comparisons test was used, as indicated in the corresponding figure legends. A p-value <0.05 was considered statistically significant.

## RESULTS

### Most G>A variants are splice-associated or occur near splice junctions

Pathogenic (P) and likely pathogenic (LP) G>A variants are attractive candidates for ADAR-mediated SDRE, however, their genomic location may influence both editing efficiency and pre-mRNA splicing. Analysis of all P/LP exonic and intronic G>A variants in ClinVar (n=40,552) revealed that 39% are possibly splice-associated, including canonical or near-canonical splice-site variants (+1 to +5 and -1 to -5), intronic splice region variants, and exonic variants predicted to affect splicing (SpliceAI score ≥0.8) (Figure 1B). Variant proximity to exon-intron boundaries may also influence SDRE, as gRNA binding across these regions has the potential to interfere with splice-site recognition. Over 39% of P/LP intronic and exonic G>A variants are located on the first or last base of the exon or one base away from this location (Figure 1C). Cumulatively, 63% were located within 30 nt of an exon-intron boundary (defined as the first or last nucleotide of an exon), 76% within 50 nt, and 89% within 100 nt (Figure 1D), indicating that a substantial proportion of clinically relevant variants fall within the binding regions of symmetric 60-, 100-, and 200-nt gRNAs respectively. These results show that the majority of G>A variants are either splice variants themselves and/or fall close to a splice boundary.

Although G>A variants are the most direct targets for ADAR-mediated SDRE, some non-G>A variants may also be amenable to editing by targeting nearby adenosines within splice-regulatory elements. Deep intronic variants illustrate this concept: they account for 0.5% of pathogenic/likely pathogenic ClinVar variants, yet only 18% of these are G>A substitutions. Because the therapeutic objective for pseudoexon-causing deep intronic variants is to prevent pseudoexon inclusion rather than correct the causative nucleotide itself, adenosines located within splice-critical regions may serve as alternative editing targets. This strategy could substantially expand the range of deep intronic variants amenable to ADAR-based SDRE and, in some cases, enable a single editing target to suppress pseudoexon inclusion caused by multiple pathogenic variants that generate the same or overlapping pseudoexons.

### Pre-mRNA and mature mRNA can be selectively edited using transcript-specific gRNAs

Exonic variants located near exon-intron boundaries are present in both unspliced pre-mRNA and mature mRNA, and we therefore asked whether gRNA design could be used to selectively direct ADAR-mediated editing toward a specific transcript type. To study the ability of an exogenous gRNA to harness the ADAR enzyme and selectively edit either the pre-mRNA or mRNA transcript of a gene, we first selected three separate adenosine targets for deamination, all at the beginning or end of exons (*ABCA4* c.4634G>A, *ABCA4* c.1556G>A, *PDE6B* c.1832A wildtype nucleotide). The *ABCA4* c.4634G>A (last base of exon 31) is a variant of uncertain significance (VUS) that has been shown to have no effect on splicing using a midigene approach in HEK293T cells (45) and *ABCA4* c.1556G>A (second base of exon 12) is a VUS and is also predicted to have no effect on splicing (SpliceAI score of 0.11) (49). *PDE6B* c.1832A is a wild-type adenosine located at the final nucleotide of exon 11, a position that is a guanosine in the majority of human exons (9.8% versus 80.4%, respectively) (50). We chose these nucleotides to be informative targets because symmetric gRNAs designed for each type of RNA transcript (pre-mRNA and mRNA) would be only half perfectly-complementary to the other target RNA transcript and would lack the double stranded RNA in the vicinity of the target that allows for the ADAR enzyme to efficiently bind and subsequently deaminate (Figure 2A) (51–53). To confirm that the three adenosine targets indeed do not affect splicing, we initially transfected previously reported midigene-containing splice vectors harboring either the *ABCA4* c.4634G>A variant or its corresponding WT sequence (45), as well as newly generated minigene-containing splice vectors harboring *ABCA4* c.1556G>A (or its corresponding WT sequence) and the wildtype *PDE6B* c.1832A region (Figure 2B) into HeLa cells. None of the constructs showed any effect on the expected canonical splicing product (Supplementary Figure 1). These results show that the three chosen target adenosines are optimal for comparison of selectively edited pre/post-spliced transcripts via ADAR-mediated SDRE with gRNAs.

**Figure 2.**
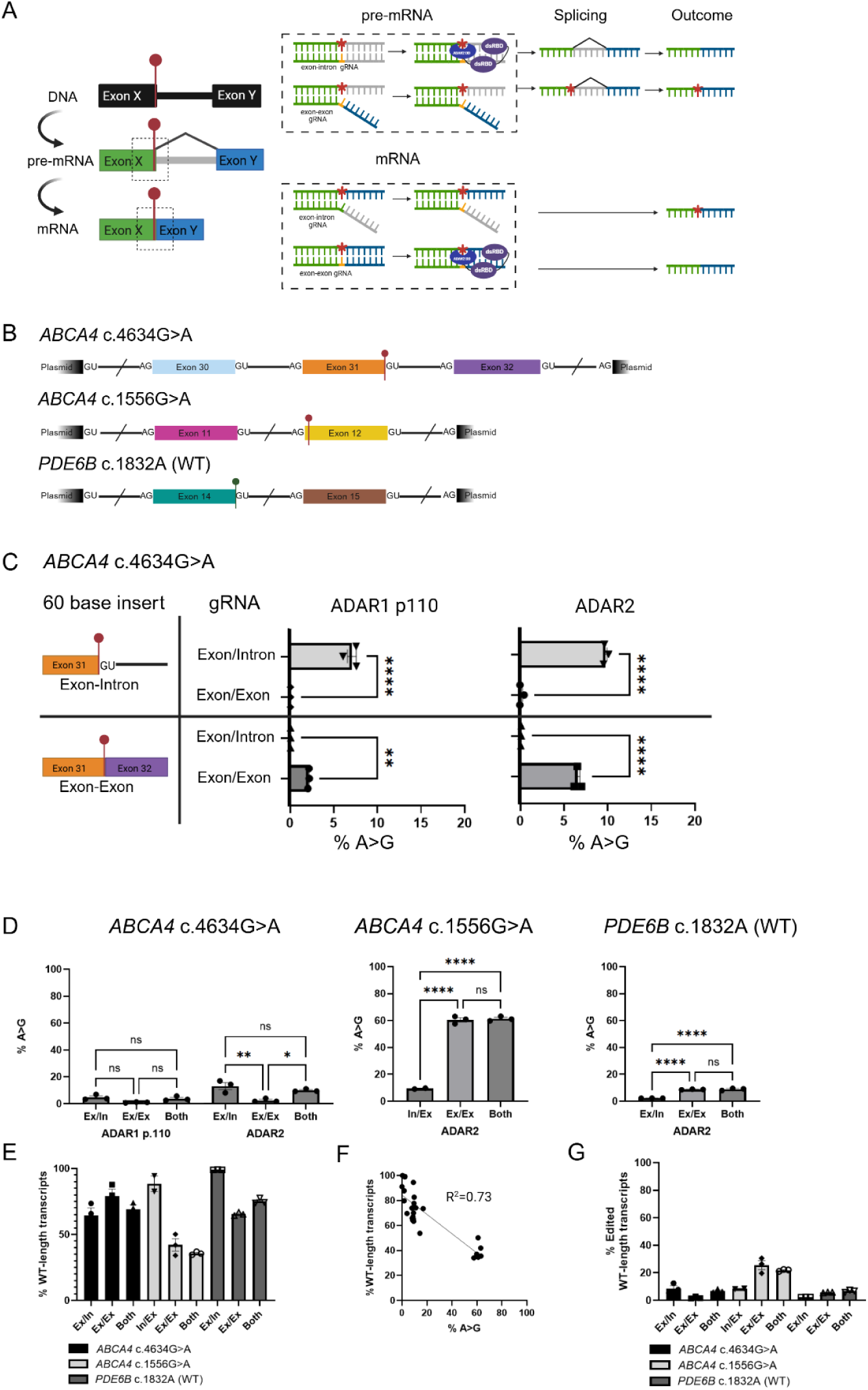
Transcript-specific SDRE enables selective targeting of pre-mRNA and mature mRNA but can induce splice perturbation. (A) Schematic representation of transcript-specific gRNAs designed to target either pre-mRNA or mature mRNA. An exon-intron gRNA directs ADAR-mediated deamination of a terminal exonic adenosine within the pre-mRNA prior to splicing, whereas an exon-exon gRNA directs editing of the corresponding mature mRNA after splicing. This design enables selective targeting of each transcript species with minimal cross-editing of the alternative transcript. (B) Schematic representation of mini/midigene splice constructs containing exonic adenosine targets positioned near exon boundaries: *ABCA4* c.4634G>A (last nucleotide of exon), *ABCA4* c.1556G>A (second nucleotide of exon), and *PDE6B* c.1832A (last nucleotide of exon). Markers above the transcript indicate target adenosines, slashes indicate shortened intronic regions, and GU and AG denote splice donor and splice acceptor sites, respectively. (C) A-to-G editing levels mediated by exon/intron (ex/in) or exon/exon (ex/ex) gRNAs in ADAR1 p110- and ADAR2-overexpressing HeLa cells targeting the *ABCA4* c.4634G>A adenosine in minimal/no-splicing constructs containing 60 nucleotides of either the exon-intron or exon-exon sequence surrounding the target site. (D) A-to-G editing levels mediated by transcript-specific gRNAs targeting splice-proximal exonic adenosines. *ABCA4* c.4634G>A was assessed in both ADAR1 p110- and ADAR2-overexpressing HeLa cells while *ABCA4* c.1556G>A and *PDE6B* c.1832A were assessed in ADAR2-overexpressing HeLa cells. (E) Percentage of total transcripts exhibiting correct wild-type (WT) splicing for each target and gRNA type in ADAR2-overexpressing HeLa cells. (F) Correlation between A-to-G editing levels and the percentage of correctly spliced wild-type (WT)-length transcripts (Pearson’s r = −0.86, R² = 0.73, n=28). (G) Percentage of total transcripts that both underwent A-to-G editing and exhibit correct wild-type (WT) splicing for each target and gRNA type in ADAR2-overexpressing HeLa cells. (C-E,G) Each data point represents an independent biological replicate; bars indicate mean ± SEM (n = 3, except n = 2 for the in/ex gRNA targeting *ABCA4* c.1556G>A in panels D, E, and G). Statistical significance was determined using one-way ANOVA with Tukey’s multiple comparisons test. ns, not significant; * *P* < 0.05, ** *P* < 0.01, and **** *P* < 0.0001. *Created in BioRender. Schneider, N.* (*2026*) https://BioRender.com/84yvecq

Before comparing editing efficiencies between pre-mRNA and mature mRNA targets, we first sought to determine whether a symmetric transcript-specific gRNA could meaningfully edit its non-target transcript type. In order to assess these editing efficiencies independent of the effects of splicing and its timing, we minimized these variables as much as possible by designing two constructs that contained only the 60bp target sequence of the exon-intron (pre-mRNA) or exon-exon (mRNA) and cloned each one into an expression construct. In both ADAR1 p110- and ADAR2-overexpressing cells, each transcript-specific gRNA preferentially edited its fully complementary target construct while showing minimal to no activity on the alternate transcript sequence (Figure 2C). Hereafter, symmetric gRNAs targeting the end of an exon and therefore complementary to exon-intron junctions are referred to as “ex/in gRNAs”, those targeting the beginning of an exon and therefore intron-exon junctions are referred to as “in/ex gRNAs”, and gRNAs complementary to exon-exon junctions are referred to as “ex/ex gRNAs.” In ADAR1 p110-overexpressing cells, the ex/in gRNA (v21.1, 5’−29-C-30, where the numbers denote the number of complementary nucleotides flanking the target site and "C" denotes the single orphan cytidine positioned opposite the target adenosine) edited the exon-intron construct at 7.1% while the ex/ex gRNA (v.21.2, 5’−29-C-30) produced no detectable editing of the exon-intron construct (n=3, p <0.0001) (Figure 2C). Conversely, the ex/ex gRNA (v.21.2) edited the exon-exon construct at 2.1%, but the ex/in gRNA (v21.1) showed only 0.1% editing on the exon-exon construct (n=3, *P* = 0.002) (Figure 2C). A similar pattern was observed in ADAR2-overexpressing cells, where the ex/in gRNA (v21.1) edited the exon-intron construct at 9.7% while the ex/ex gRNA (v21.2) showed 0.1% editing of the same construct (n=3, *P* < 0.0001), and the ex/ex gRNA (v21.2) preferentially edited the exon-exon construct with 6.6% efficiency while the ex/in gRNA (v21.1) showed only 0.1% editing of this construct (n=3, *P* < 0.0001) (Figure 2C). Together, these findings demonstrate that for these targets and gRNAs, RNA editing required more than half complementarity between the gRNA and transcript sequences, allowing selective targeting of either pre-mRNA or mature mRNA sequences.

As little is known about using gRNA to selectively and exclusively edit unspliced and spliced coding RNA, we wanted to see if it is possible to edit these separately, if there is a difference between the editing levels, and if it may be possible to leverage the different transcripts for an additive effect by using two different kinds of gRNA at once. For each variant described in this section, two chemically modified symmetric 60-nt gRNAs were designed: an ex/in or in/ex gRNA targeting unspliced pre-mRNA and an ex/ex gRNA targeting spliced mRNA, all target RNA derived from a mutant splice construct. RNA editing was measured from WT-length transcripts that underwent proper splicing. For *ABCA4* c.4634G>A, in both ADAR1 p110- and ADAR2-overexpressing cells, the ex/in gRNA preferentially edited the pre-mRNA transcript compared with editing of mature mRNA by the ex/ex gRNA (Figure 2D). In ADAR1 p110-overexpressing cells, the ex/in gRNA (v21.1) resulted in 4.7% pre-mRNA editing compared with 1.0% mature mRNA editing by the ex/ex gRNA (v21.2), although this difference did not reach statistical significance (n=3, *P* = 0.053). A stronger effect was observed in ADAR2-overexpressing cells, where the ex/in gRNA (v21.1) resulted in 13.2% pre-mRNA editing compared with 2.5% mature mRNA editing by the ex/ex gRNA (v21.2) (n=3, *P* = 0.008). A different editing preference emerged for the *ABCA4* c.1556G>A and *PDE6B* c.1832A targets. As the retina is one of the few tissues in which ADAR2 is most strongly expressed (54, 55), we compared editing levels for these target adenosines for each gRNA only in ADAR2-overexpressing HeLa cells. Both showed that the ex/ex gRNA preferentially edited the mRNA transcript compared with editing of pre-mRNA by the ex/in or in/ex gRNAs (Figure 2D). For *ABCA4* c.1556G>A, the ex/ex gRNA (v50.2, 5’−29-C-30) edited the mRNA transcript at a sixfold higher rate than the in/ex gRNA (v50.1, 5’−29-C-30) (60.5%, n=3 and 9.5%, n=2, respectively, *P* < 0.0001) (Figure 2D). *PDE6B* also showed this preference pattern, albeit with lower editing levels, with the ex/ex gRNA (v70.2, 5’−29-C-30) editing the mRNA transcript at a fourfold higher rate than the ex/in gRNA (v70.1, 5’−29-C-30) (8.8%, n=3 and 2.3%, n=3, respectively, *P* < 0.0001) (Figure 2D). Cotransfection of both gRNAs resulted in detectable editing for all target adenosines mentioned but did not appear to produce an additive increase in editing efficiency, as none of the editing levels differed significantly from the higher-efficiency gRNA (Figure 2D). These results suggest that it is possible to exclusively target a transcript type (pre-mRNA or mRNA) for SDRE, that certain transcripts may be preferentially edited depending on the sequence context, and that combining two gRNA approaches may not lead to increased total editing but also do not seem to compete in a detrimental manner.

### Correctly spliced edited transcripts remain achievable despite splice perturbation by gRNAs

Because the three above-mentioned target adenosines are located in close proximity to splice junctions, gRNA binding has the potential to interfere with splice-site recognition and nearby ESE elements. We therefore assessed the impact of transcript-specific gRNAs on splicing in parallel with editing efficiency. Across all three targets, gRNA introduction was accompanied by varying degrees of mis-splicing, regardless of gRNA type (Figure 2E).

Notably, a relationship emerged between editing efficiency and splice perturbation. Across all transcript-specific gRNA experiments, higher editing levels were associated with greater disruption of normal splicing, resulting in a strong inverse correlation between editing efficiency and the proportion of correctly spliced transcripts (Pearson’s r = −0.86, R² = 0.73; Figure 2F). Though this was the case, the final amounts of edited wildtype-length transcripts could reach 25.6% of total final transcripts (Figure 2G). These findings suggest that while gRNA binding and ADAR-mediated editing near splice junctions may increase the likelihood of splice perturbation, such effects do not necessarily preclude the generation of substantial levels of correctly spliced edited transcripts.

### SDRE targeting of a splice-critical adenosine can reduce pseudoexon inclusion

Having demonstrated that pre-mRNA can be selectively targeted and edited by exogenous gRNAs in our cellular system, together with extensive prior evidence showing that intronic regions are frequent endogenous substrates of ADAR-mediated editing in humans (17, 56), we next explored whether SDRE could be harnessed as a targeted approach for splice modulation within deep intronic regions. Digital droplet PCR (ddPCR) results showed that for three *ABCA4* midigene constructs, the first two of which have been previously reported (46), harboring either the WT sequence, c.5196+1137G>A, or c.5196+1137G>T (Figure 3A), revealed marked differences in pseudoexon inclusion following transfection into HeLa cells (Supplementary figure 2A). The pathogenic c.5196+1137G>A variant produced minor pseudoexon inclusion (6.7%, n=2) and the c.5196+1137G>T variant resulted in substantially higher pseudoexon incorporation (46.7%, n=2) (Supplementary Figure 2A). A similar pattern was observed in 661W cells, where the splice defect associated with c.5196+1137G>T appeared even more pronounced (Supplementary Figure 2B).

**Figure 3.**
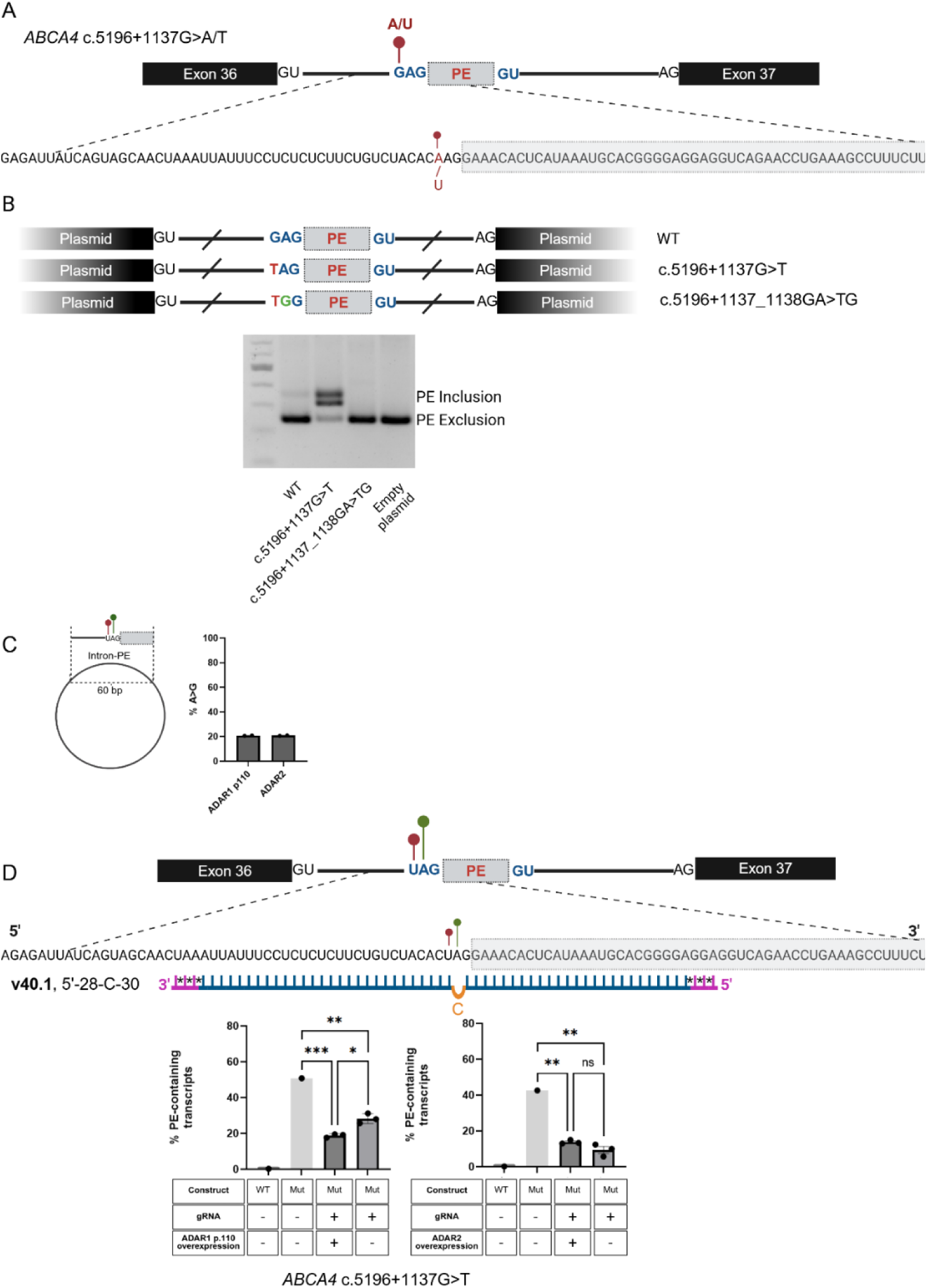
SDRE targeting of a splice-critical adenosine reduces pseudoexon inclusion in ADAR1 p110-overexpressing cells. (A) Schematic representation of the *ABCA4* c.5196+1137G>A and c.5196+1137G>T deep intronic variants and the resulting 73 base pseudoexon (PE). A red marker above the transcript indicates the RNA-level consequence of the variant (G>A/U). GU and AG denote splice donor and splice acceptor sites, respectively. (B) Top: Schematic representation of minigene splice constructs containing either the WT sequence, the experimentally assessed c.5196+1137G>T variant, or the c.5196+1137_1138GA>TG sequence mimicking complete A-to-G editing of the target adenosine. Bottom: Agarose gel analysis of PCR products showing pseudoexon (PE) inclusion (319 bp) and/or exclusion (246 bp). The upper band migrating approximately 20 nt above the pseudoexon-containing product was identified by gel excision and sequencing as a PCR heteroduplex. (C) Left: Schematic representation of a minimal/no-splicing construct containing 60 nucleotides of intronic sequence surrounding the target site. Right: A-to-G editing levels mediated by gRNA v40.1 in ADAR1 p110- and ADAR2-overexpressing HeLa cells targeting the splice acceptor adenosine (ABCA4 c.5196+1138A) in the minimal/no-splicing construct. Bars indicate mean ± SEM (n = 2). (D) Top: Schematic representation of the *ABCA4* c.5196+1137G>T deep intronic variant (red marker) and c.5196+1138 adenosine target (green marker), resulting pseudoexon (PE), and gRNA v40.1 design. Pink nucleotides indicate 2′-O-methyl modifications, asterisks indicate phosphorothioate linkages, blue nucleotides indicate RNA bases, and the orange nucleotide indicates the orphan cytosine-to-adenosine mismatch opposite the target adenosine. Bottom: ddPCR quantification of the percentage of PE-retaining transcripts following treatment with gRNA v40.1 in ADAR1 p110- and ADAR2-overexpressing or uninduced HeLa cells. Bars indicate mean ± SEM (n = 3, except n = 1 for gRNA-untreated WT and mutant samples). Statistical significance was determined using one-way ANOVA with Tukey’s multiple comparisons test for predefined comparisons. ns, not significant; * *P* < 0.05, ** *P* < 0.01, *** *P* < 0.001. *Created in BioRender. Schneider, N.* (*2026*) https://BioRender.com/adv63d8

We hypothesized that if the c.5196+1137G>T variant strengthens the aberrant splice acceptor region by generating a favorable -3 acceptor context, we could target a close-by adenosine for RNA editing to weaken or ablate this splice acceptor motif. We therefore selected the 3’ neighboring adenosine (*ABCA4* c.5196+1138A) as the target for ADAR-mediated editing, reasoning that A-to-I editing at this position could disrupt splice acceptor recognition and thereby suppress pseudoexon inclusion, since in 99.7% of splice acceptor sites, there is an adenosine in this location (50). To test this hypothesis, we simulated complete A>G editing at this position by generating minigene splice constructs containing either the WT sequence, the c.5196+1137G>T variant, or the c.5196+1137_1138GA>TG sequence mimicking total A>G editing of the neighboring adenosine (Figure 3B). Gel electrophoresis analysis demonstrated clear pseudoexon inclusion in the c.5196+1137G>T mutant construct, whereas the c.5196+1137_1138GA>TG editing-simulation construct showed no detectable pseudoexon inclusion (Figure 3B). An additional, slower-migrating band observed in some samples was identified by gel excision and Sanger sequencing as a heteroduplex formed during PCR and therefore does not represent a distinct splice isoform. Together, these findings demonstrate that the splice acceptor adenosine adjacent to c.5196+1137G>T represents a viable SDRE target and that disruption of this -2 splice acceptor position may prevent aberrant pseudoexon recognition within the deep intron.

To assess the intrinsic editability of the target sequence independent of ongoing splicing dynamics by a 59-mer chemically modified symmetric gRNA (v40.1, 5’−28-C-30), we generated a minimal/no splicing construct containing the 60bp target region harboring the *ABCA4* c.5196+1137G>T variant cloned into an expression vector (Figure 3C). This system enabled direct quantification of RNA editing without the effects of transcript processing. Stable HeLa cell-lines overexpressing either ADAR1 p110 or ADAR2 were transfected with both the target construct and the corresponding gRNA (v40.1). Similar editing efficiencies were observed in both cellular systems, with ADAR1 p110- and ADAR2-overexpressing cells reaching 20.5% and 20.9% editing, respectively (Figure 3C). These findings demonstrate that the designed gRNA can recruit both ADAR1 p110 and ADAR2 to the relevant 60 bases of the deep intronic target sequence for targeted editing.

Due to the deep intronic location of this target nucleotide, direct quantification of A-to-G editing was not feasible, as the target nucleoside is spliced out from both the wildtype transcript and the pseudoexon-containing transcript. We therefore evaluated the ADAR enzyme’s effect indirectly by quantifying changes in pseudoexon inclusion using ddPCR. ddPCR probes were designed to selectively detect either the correctly spliced transcript by binding to the exon 36/exon 37 junction or the aberrantly spliced transcript containing the pseudoexon. Midigene constructs and gRNA (v40.1) were transfected into ADAR1 p110- and ADAR2-overexpressing HeLa cells or uninduced HeLa cells expressing baseline endogenous ADAR levels. ddPCR analysis revealed that all conditions in which gRNA v40.1 was introduced resulted in a significant reduction in pseudoexon inclusion compared with the mutant control (Figure 3D). In addition, ADAR1 p110 overexpression further reduced pseudoexon inclusion compared with endogenous ADAR levels (18.8% versus 28.3%, respectively; n=3, *P* = 0.017; Figure 3D). In contrast, ADAR2 overexpression did not result in a significant difference in pseudoexon inclusion between conditions (Figure 3D). These findings suggest that for this target, increased ADAR1 p110 expression together with gRNA-mediated recruitment to the target transcript promotes restoration of correct splicing within this cellular system, potentially through editing-mediated disruption of the aberrant splice acceptor.

### Near-canonical splice-site variants may be especially vulnerable to guide-induced splice disruption

To further explore the application of ADAR-mediated SDRE on pre-mRNA targets, we selected the previously characterized pathogenic near-canonical splice-site variant *ABCA4* c.5714+5G>A, which is located in close proximity to the exon 40/intron 40 junction (Figure 4A) and has been reported to cause partial exon 40 skipping (45). Previously reported midigene splice constructs harboring either the WT or mutant sequence were used to evaluate the effects of targeted editing on splicing outcomes (45).

**Figure 4.**
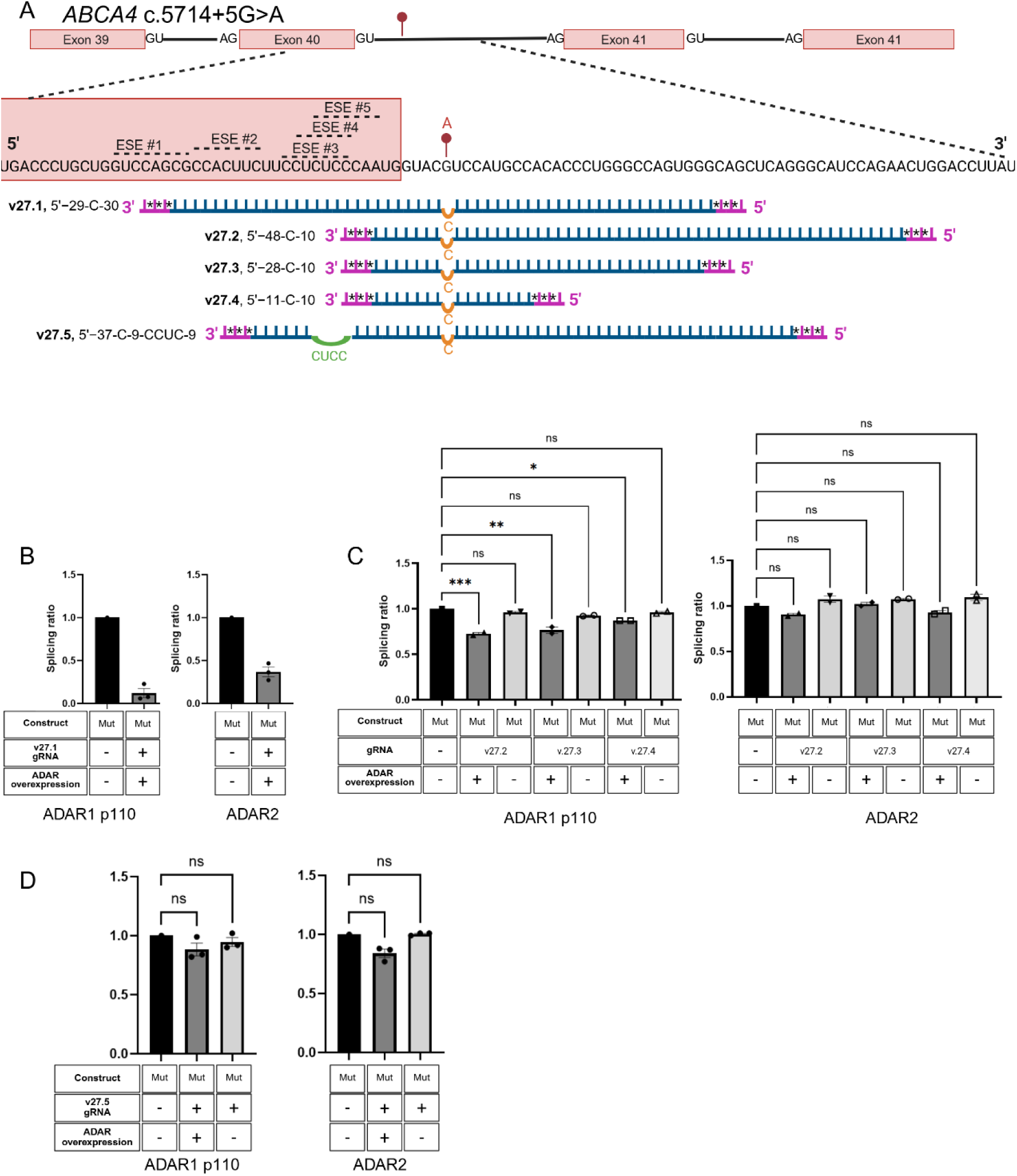
Optimization of gRNAs targeting a canonical splice-site mutation reduces gRNA-mediated splice perturbation but does not rescue mutation-induced splicing defects. (A) Schematic representation of the *ABCA4* c.5714+5G>A splice donor variant and the gRNA designs evaluated. The red marker indicates the mutant adenosine targeted for editing. Dashed lines denote predicted ESE motifs within the gRNA binding regions. For each gRNA, pink nucleotides indicate 2′-O-methyl modifications, asterisks indicate phosphorothioate linkages, blue nucleotides indicate RNA bases, green nucleotides indicate a four nt bulge, and the orange nucleotide indicates the orphan cytosine opposite the target adenosine. (B) Comparison of splicing ratios in uninduced HeLa cells and ADAR1 p110- or ADAR2-overexpressing HeLa cells following treatment with symmetrical gRNA v27.1. (C) Comparison of splicing ratios in mutant construct-transfected untreated uninduced HeLa cells, gRNA-treated uninduced HeLa cells, and gRNA-treated ADAR1 p110- or ADAR2-overexpressing HeLa cells using asymmetrical gRNAs v27.2, v27.3, or v27.4. Splicing ratio was calculated as the percentage of wild-type (WT) transcripts in each experimental condition relative to the untreated mutant control (D) Comparison of splicing ratios in mutant construct-transfected untreated and uninduced HeLa cells, gRNA-treated uninduced HeLa cells, and gRNA-treated ADAR1 p110- or ADAR2-overexpressing HeLa cells using the asymmetrical and loop-containing gRNA v27.5. (B–D) Splicing ratio was calculated as the percentage of wild-type-length (WT) transcripts relative to the untreated mutant control. Each data point represents an independent biological replicate; bars indicate mean ± SEM. Sample sizes were n = 3 (B), n = 2 (C), and n=3 (D); the untreated mutant control in uninduced cells was n = 1. Statistical significance was determined using one-way ANOVA with Tukey’s multiple comparisons test; not all comparisons are shown. ns, not significant; * *P* < 0.05, ** *P* < 0.01, *** *P* < 0.001. *Created in BioRender. Schneider, N.* (*2026*) https://BioRender.com/j5hu3gy

We initially designed a symmetrical, chemically modified 60-mer gRNA (v27.1, 5’−29-C-30) spanning the exon-intron boundary. Analysis with ESEfinder 3.0 predicted that the exonic portion of this gRNA could completely overlap four and partially overlap one ESE motifs (ESEs #1-5, Figure 4A). Following transfection of the mutant splice construct and gRNA v27.1 into ADAR1 p110- and ADAR2-overexpressing HeLa cells, capillary electrophoresis analysis revealed a marked decrease in properly spliced transcripts for mutant constructs when gRNA was introduced in conjunction with ADAR1 p110 and ADAR2 overexpression (Figure 4B), indicating that this gRNA design may interfere with normal splice regulation in conjunction with ADAR.

To reduce overlap with predicted ESE motifs, we next designed three asymmetric gRNAs with shortened exonic regions (Figure 4A). All three guides shared the same 5′ exonic sequence up to the C-to-A mismatch opposite the target adenosine but differed in the length of the regions binding to the intron. The three asymmetric gRNA designs, a 59-mer (v27.2, 5’−48-C-10), 39-mer (v27.3, 5’−28-C-10), and a 22-mer (v27.4, 5’−11-C-10) were designed to reduce overlap with the five predicted ESE motifs, resulting in only partial overlap with three motifs (ESEs #3,#4,#5). Capillary electrophoresis analysis showed that none of the three asymmetric gRNAs significantly altered splicing relative to the mutant construct alone (Figure 4C) when ADAR overexpression was not induced. In contrast, ADAR1 p110-overexpressing cells exhibited a reduction in correctly spliced transcripts for all three gRNAs, whereas no significant change in splicing was observed in ADAR2-overexpressing cells. These findings suggest that shortening the exonic portion of the gRNA reduced splice perturbation with endogenous ADAR and ADAR2 overexpression but still induced more mis-splicing in ADAR1 p110-overexpressing cells, and therefore no evidence of editing or splice rescue was present for all conditions.

Finally, we designed a 60-mer asymmetric gRNA (v27.5, 5’−37-C-9-CCUC-9) incorporating a four-nucleotide internal loop opposite the sequence shared by ESE motifs #3,4,5 (“CUCC”). This design, v27.5, allowed extension of the exonic portion of the gRNA from the previously mentioned 59, 39, and 22-mers while minimizing direct masking of the predicted splice-regulatory elements (Figure 4A). The 5’ portion of the gRNA was still shortened from the symmetric v27.1 from 30 nts to 22 nts to avoid ESEs #1 and 2. Cotransfection of the mutant splice construct and the v27.5 gRNA into uninduced and ADAR1 p110 or ADAR2-overexpressing HeLa cells revealed no significant difference in splicing relative to the mutant construct alone, either in the presence or absence of ADAR1 p110 or ADAR2 overexpression (Figure 4D). In contrast to the previously tested designs for this mutation, incorporation of the internal loop eliminated the splice-disruptive effects associated with gRNA binding for both ADAR1 p110- and ADAR2-overexpressing HeLa cells. However, no improvement in exon 40 inclusion was observed under these conditions. Together, these findings indicate that splice perturbation can be minimized through careful gRNA design, although editing of the *ABCA4* c.5714+5G>A target for the improvement of splicing outcomes in this experimental system was not feasible and may point to near-canonical/canonical intronic splice-site mutations being especially difficult targets for ADAR-mediated SDRE.

### ADAR-mediated RNA editing of exonic splice variants can modulate aberrant splicing

The application of site-directed ADAR-mediated RNA editing to exonic splice mutations remains relatively unexplored, with only a limited number of studies investigating its effects on splicing outcomes (25). Such variants present unique challenges for SDRE, as their proximity to exon-intron boundaries may influence both editing efficiency and splice-site recognition. In addition, exonic splice mutations may reside within ESE motifs, raising the possibility that gRNA binding itself could perturb normal splicing and promote exon skipping. To investigate the feasibility of editing exonic splice mutations, we selected two disease-associated variants located at the final nucleotide of an exon: *CDH23* c.5712G>A and *BBS1* c.479G>A.

*CDH23* c.5712G>A, a synonymous variant associated with Usher syndrome type 1D, has previously been shown in a splice assay to induce skipping of exon 42 in COS-7 cells (57). However, in-silico prediction tools have yielded conflicting predictions regarding its splicing consequence, suggesting exon skipping, intron retention, or activation of cryptic splice sites (57, 58). To further characterize the splicing effects of *CDH23* c.5712G>A, we generated minigene constructs containing either the mutant or wild-type exon 42 sequence and transfected them into HeLa cells (Figure 5A). Capillary electrophoresis analysis demonstrated the production of multiple transcript isoforms, including the highest proportion of transcripts exhibiting intron 42 retention followed by exon 42 skipping, alongside a smaller proportion of correctly spliced transcripts (Figures 5B, C).

**Figure 5:**
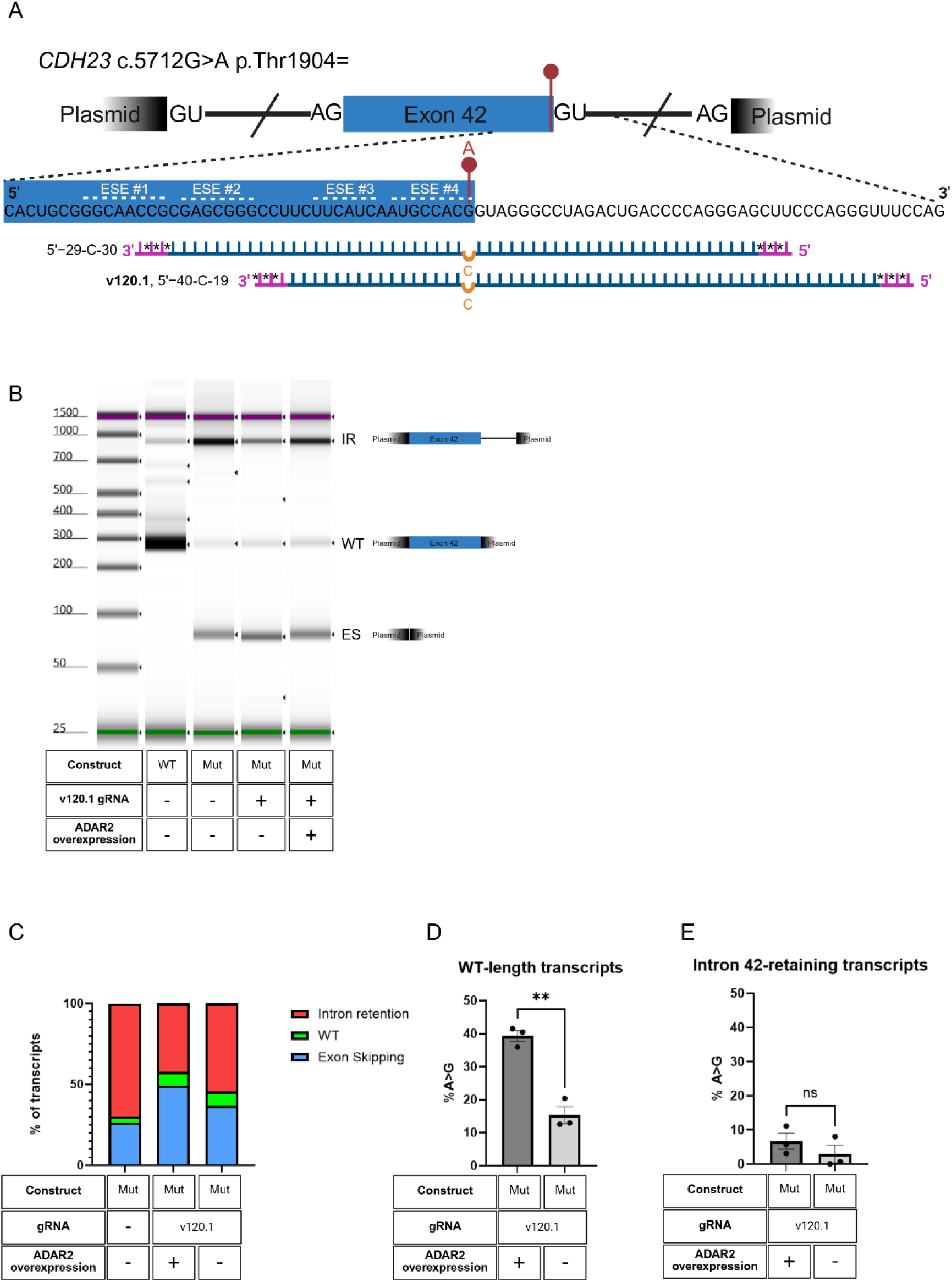
Asymmetric gRNA introduction improves *CDH23* splice-mutation outcomes with distinct splicing profiles in the presence or absence of ADAR2 overexpression. (A) Schematic representation of the *CDH23* c.5712G>A exonic splice variant in a minigene splice construct and the gRNA designs evaluated. Red marker above the transcript indicates the target adenosine, slashes indicate shortened intronic regions, and GU and AG denote splice donor and splice acceptor sites, respectively. Dashed lines denote predicted exonic splicing enhancer (ESE) motifs within the gRNA binding regions. For each gRNA, pink nucleotides indicate 2′-O-methyl modifications, asterisks indicate phosphorothioate linkages, blue nucleotides indicate RNA bases, and the orange nucleotide indicates the orphan cytosine opposite the target adenosine. (B) Capillary electrophoresis analysis comparing splicing profiles of WT splice construct-transfected uninduced HeLa cells, mutant splice construct-transfected untreated uninduced HeLa cells, mutant splice construct-transfected and gRNA v120.1-treated ADAR2-overexpressing HeLa cells, and mutant splice construct-transfected and gRNA v120.1-treated uninduced HeLa cells. IR, intron retention; WT, wild-type length transcripts; ES, exon skipping. (C) Percentage of intron-retaining (IR), exon-skipped (ES), and wild-type length (WT) transcripts in mutant construct-transfected untreated uninduced HeLa cells, gRNA v120.1-treated ADAR2-overexpressing HeLa cells, and gRNA v120.1-treated uninduced HeLa cells. (D) A-to-G editing levels within WT-length transcripts in ADAR2-overexpressing and uninduced HeLa cells. (E) A-to-G editing levels within intron 42-retaining transcripts in ADAR2-overexpressing and uninduced HeLa cells. (D-E) Each data point represents an independent biological replicate; bars indicate mean ± SEM (n = 3). Statistical significance was determined using Welch’s two-tailed unpaired *t*-test. ns, not significant, ** *P* < 0.01. *Created in BioRender. Schneider, N.* (*2026*) https://BioRender.com/qt3gr7q

Because gRNA binding across exonic sequences may interfere with splice-regulatory elements, we used ESEfinder 3.0 to map ESE motifs surrounding the variant (43, 44). This analysis was used to identify gRNA designs that minimized overlap with predicted ESEs and thereby reduced the likelihood of gRNA-mediated splice disruption. Four motifs were predicted to fully or partially exist within the 30 bases upstream of the mutation, and therefore an asymmetric 60-mer gRNA (v120.1, 5’−40-C-19) was determined to be the best candidate, as it would bind to only ESE #3 and 4 while retaining the 60-base length, and binds to both exon 42 and intron 42 in the pre-mRNA (Figure 5A).

Minigene constructs harboring either the WT or *CDH23* c.5712G>A mutant sequence were cotransfected with the v120.1 gRNA into either uninduced HeLa cells or ADAR2-overexpressing HeLa cells. No detectable splicing abnormalities were observed following transfection of the WT construct together with the gRNA, suggesting that gRNA binding alone does not induce a splicing defect in this context (Supplementary Figure 3). Introduction of the gRNA into cells transfected with the mutant construct resulted in an approximately two-fold increase in correctly spliced WT-length transcripts, from 4.0% in cells receiving the mutant construct alone to 8.4% and 8.9% in uninduced and ADAR2-overexpressing HeLa cells, respectively (Figure 5B,C). This increase in WT-length transcripts was accompanied by a shift in the distribution of aberrantly spliced isoforms, characterized by increased exon 42 skipping and reduced intron 42 retention. Notably, the magnitude of these changes differed between the two cellular backgrounds, with ADAR2-overexpressing cells exhibiting a smaller increase in exon skipping and a greater increase in intron retention (Figure 5C).

Analysis of editing levels revealed substantial editing of the target adenosine within correctly spliced WT-length transcripts, reaching 42.1% in ADAR2-overexpressing cells compared with 15.8% in uninduced HeLa cells (Figure 5D). Due to the location of the target adenosine and gRNA design, it is reasonable to assume the editing for these WT-length transcripts occurs before splicing. Additionally, transcripts retaining intron 42 exhibited only low levels of editing in both uninduced and ADAR2-induced HeLa cells with no significant difference between them (5.5% and 3.1%, respectively) (Figure 5E). Together, these findings indicate that gRNA introduction increased the abundance of correctly spliced WT-length transcripts, while ADAR2 overexpression was associated with substantially higher editing levels within these transcripts. In addition, ADAR2-overexpressing cells exhibited increased intron retention following gRNA transfection, although these intron-retaining transcripts showed only low levels of editing not statistically significantly different from endogenous ADAR-expressing cells. These observations suggest that ADAR2 expression influences both editing efficiency and splicing outcomes within this system.

*BBS1* c.479G>A is a previously reported exonic splicing variant on the last base of exon 5, causing a mild form of Bardet-Biedl syndrome (59, 60), characterized by intron 5 retention or exon 5 skipping. We designed a minigene splice construct containing exons 5-7, harboring either the mentioned variant or WT sequence, and transfected each into HeLa cells (Figure 6A). Capillary electrophoresis analysis demonstrated that the mutant minigene produced multiple transcript isoforms, including the highest proportion of transcripts exhibiting exon 5 skipping followed by intron 5 retention with no visible WT transcripts (Figure 6B). This splicing pattern differs from the pattern reported in patient EBV-transformed lymphoblastoid cell lines, venous blood, and patient fibroblasts, showing higher proportions of intron inclusion and remaining WT-length bands (59, 60).

**Figure 6:**
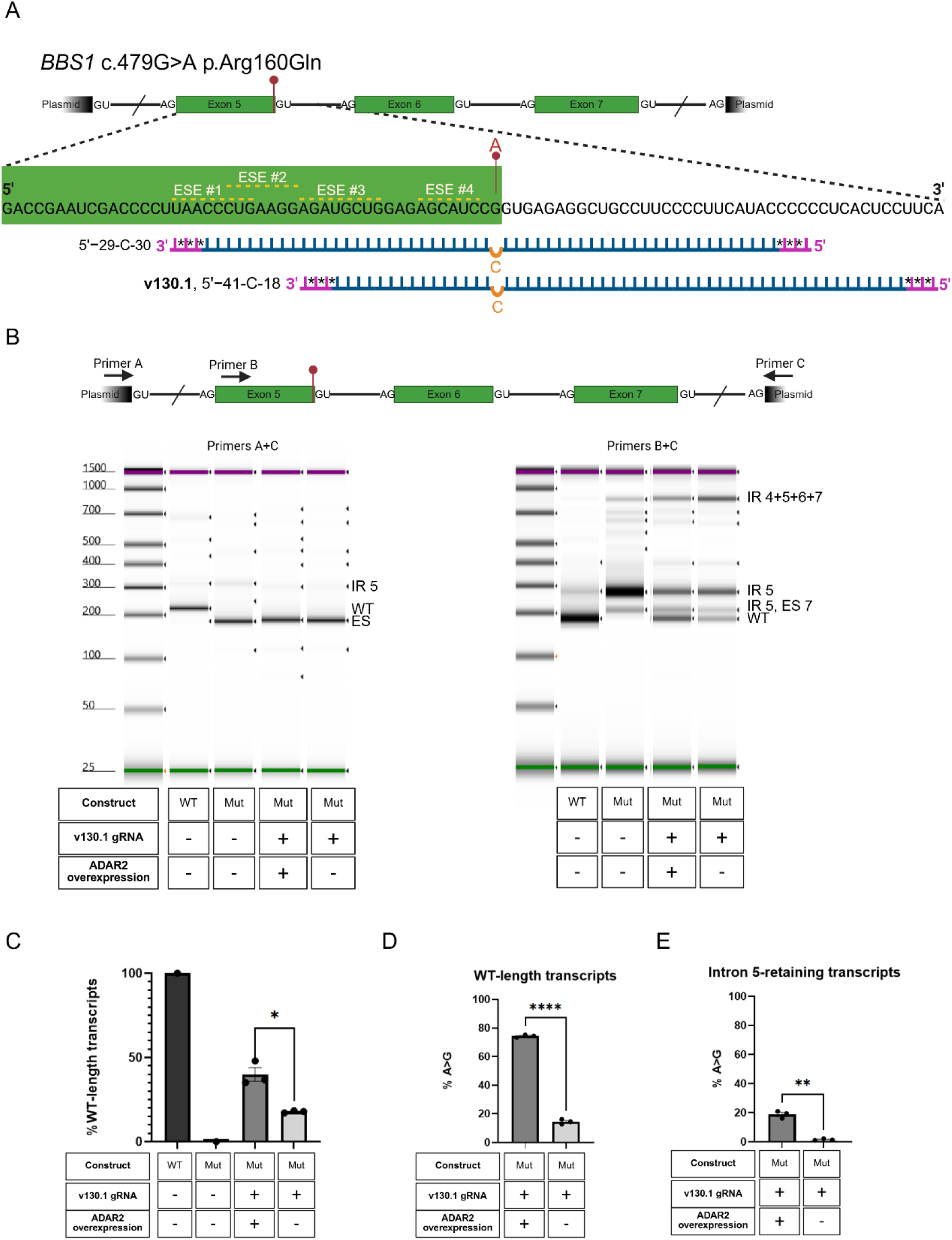
Asymmetric gRNA facilitates SDRE and reduces intron retention for *BBS1* splice mutation. (A) Schematic representation of the *BBS1* c.479G>A exonic splice variant in the minigene splice construct and the gRNA designs evaluated. The red marker indicates the target adenosine, slashes indicate shortened intronic regions, and GU and AG denote splice donor and splice acceptor sites, respectively. Dashed lines denote predicted exonic splicing enhancer (ESE) motifs within the gRNA binding regions. For each gRNA, pink nucleotides indicate 2′-O-methyl modifications, asterisks indicate phosphorothioate linkages, blue nucleotides indicate RNA bases, and the orange nucleotide indicates the orphan cytosine opposite the target adenosine. (B) Top: Schematic showing the locations of the PCR primers. Bottom: Capillary electrophoresis analysis of splicing profiles from WT or mutant splice construct-transfected uninduced HeLa cells and mutant splice construct-transfected ADAR2-overexpressing or uninduced HeLa cells treated with gRNA v130.1. Left: PCR products amplified using primers A+C. Right: PCR products amplified using primers B+C. IR 4+5+6+7, unspliced transcript retaining introns 4, 5, 6, and 7; IR 5, intron 5 retention; IR 5 ES 7, intron 5 retention with exon 7 skipping; WT, wild-type-length transcripts; ES, exon skipping. (C) Percentage of WT-length transcripts measured using primers B+C. (D) A-to-G editing levels within WT-length transcripts amplified using primers B+C. (E) A-to-G editing levels within intron 42-retaining transcripts amplified using primers B+C. (C–E) Each data point represents an independent biological replicate; bars indicate mean ± SEM. Sample sizes were n = 3 for all experimental groups, except the WT and mutant controls in (C) (n = 1). Statistical significance was determined using one-way ANOVA with Tukey’s multiple comparisons test in (C) with not all comparisons shown and Welch’s two-tailed unpaired *t*-test in (D–E). * *P* < 0.05; ** *P* < 0.01; **** *P* < 0.0001. *Created in BioRender. Schneider, N.* (*2026*) https://BioRender.com/2322zzu

Four ESE motifs were identified by ESEfinder 3.0 within 30 nucleotides of the *BBS1* c.479G>A variant on exon 5 (Figure 6A). To minimize interference with predicted splice-regulatory elements, we designed an asymmetric 60-mer gRNA spanning 18 nucleotides of exon 5 and 41 nucleotides of the adjacent intron (v130.1, 5’−41-C-18), thereby avoiding two of the predicted ESE motifs (Figure 6A). In both uninduced or ADAR2-overexpressing HeLa cells transfected with the mutant construct and gRNA, PCR on cDNA using primers spanning the first and last exons of the splice construct failed to detect WT-length transcripts due to the predominance of exon 5-skipped isoforms (Figure 6B).

To better assess any remaining correctly spliced transcripts, we performed PCR on the cDNA using primers spanning exon 5 to the terminal exon of the construct, thereby excluding exon 5-skipped isoforms from the analysis. These splicing products in ADAR2-overexpressing cells transfected with the mutant construct and v130.1 gRNA exhibited a significantly higher proportion of WT-length transcripts than uninduced cells (39.9% versus 17.8%, respectively; *P* = 0.03) (Figure 6B,C). This increase was accompanied by substantially higher editing levels within the WT-length transcript population in ADAR2-overexpressing cells compared with uninduced cells (74.34% versus 14.46%, respectively; *P* < 0.0001) (Figure 6D). Intron-retaining samples showed 4-fold less editing than their WT-length counterparts (Figure 6E).

Given the location of the target adenosine at the exon-intron boundary and the design of the gRNA, it is likely that editing occurred predominantly on the pre-mRNA prior to splicing. Although editing did not restore normal exon inclusion in this system, increased editing was associated with a positive shift in splicing outcomes characterized by reduced intron retention and enrichment of correctly spliced WT-length transcripts.

### Rational gRNA optimization increases correctly spliced edited transcripts

The interplay between ADAR-mediated SDRE and splicing observed in the preceding experiments prompted us to investigate whether the splice-disruptive effects of gRNA binding to pre-mRNA could be minimized while maintaining efficient editing. As a proof of concept for gRNA optimization, we selected the *ABCA4* c.1556G>A variant. As shown above (Figure 2D,E) gRNA v50.2 targeting the exon-exon boundary, achieved editing levels of up to 60% within correctly spliced WT-length transcripts, but this was accompanied by substantial splice disruption, with more than half of all transcripts exhibiting aberrant splicing mostly characterized by the skipping of one or more exons (Figure 2E).

Because the gRNA v50.2 spans the exon 11-exon 12 junction, we hypothesized that gRNA binding to these specific exonic regions may interfere with splice-regulatory elements and contribute to the observed mis-splicing, as several ESE motifs were predicted to be in this region (Figure 7A). To identify regions most likely to drive these effects, both ESE-related and unrelated, we performed an ASO walk across the terminal portion of exon 11 and the beginning of exon 12 in HeLa cells transfected with the minigene construct (Figure 7A). Analysis of splicing outcomes by gel electrophoresis identified five regions whose targeting was associated with pronounced splice disruption: the first three nucleotides of the target region (covered by ASO1), the final two nucleotides of exon 11 (covered by ASO5), the first six nucleotides of exon 12 including the target adenosine (covered by ASO6), a nine-nucleotide region within exon 12 (covered by ASO6-9), and the final four nucleotides of exon 12 (covered by ASO9) (Figure 7B).

**Figure 7.**
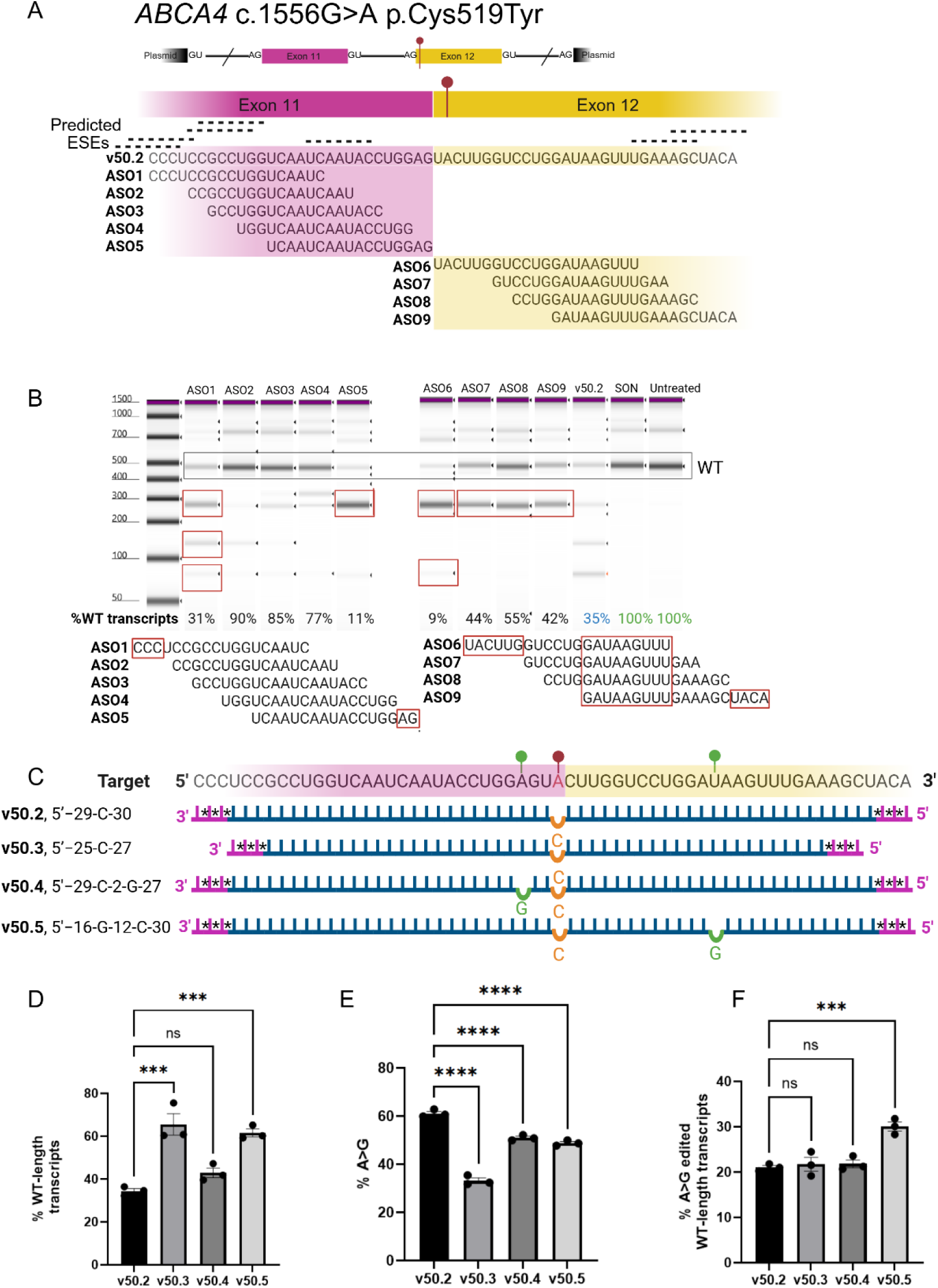
Identification of splice-critical regions by an ASO walk enables gRNA optimization to increase total edited and properly spliced transcripts. (A) Schematic representation of the *ABCA4* c.1556G>A exonic variant splice construct and the antisense oligonucleotide (ASO) RNA targets. The red marker above the transcript indicates the target adenosine for SDRE, slashes indicate shortened intronic regions, and GU and AG denote splice donor and splice acceptor sites, respectively. Dashed lines denote predicted exonic splicing enhancer (ESE) motifs within the ASO binding regions. (B) Capillary electrophoresis analysis comparing splicing patterns of mutant splice construct-transfected HeLa cells treated with ASOs #1-9, gRNA v50.2, a scrambled oligonucleotide (SON) control, or left untreated. WT, wild-type length transcript. Black box highlights WT-length transcript bands, whereas red boxes highlight aberrantly spliced transcript bands and the corresponding predicted splice-critical regions targeted by ASOs that induced mis-splicing. (C) Top: Schematic representation of the editing target region showing the *ABCA4* c.1556G>A exonic variant (red marker) and nucleotide positions selected for introduction of mismatches (green markers) based on the ASO walk. Bottom: Schematic representation of gRNAs v50.2-v50.5. Pink nucleotides indicate 2′-O-methyl modifications, asterisks indicate phosphorothioate linkages, blue nucleotides indicate RNA bases, green nucleotides indicate G-to-A mismatches introduced to reduce splice interference, and the orange nucleotide indicates the orphan C-to-A mismatch opposite the target adenosine for editing. (D) Percent wild-type length transcripts for mutant splice construct-transfected ADAR2-overexpressing HeLa cells treated with gRNAs v50.2-50.5. (E) A-to-G editing levels of *ABCA4* c.1556G>A mediated by gRNAs v50.2-v50.5 in ADAR2-overexpressing HeLa cells. (F) Percentage of total transcripts that underwent A-to-G editing and exhibit correct wild-type (WT) splicing for gRNAs v50.2-v50.5 in ADAR2-overexpressing HeLa cells. (B,D) % WT transcripts represents the percentage of WT-length transcripts relative to total detected transcripts, normalized to the untreated mutant control. (D-F) Each data point represents an independent biological replicate; bars indicate mean ± SEM (n = 3). Statistical significance was determined using one-way ANOVA with Tukey’s multiple comparisons test for predefined comparisons. ns, not significant; *** *P* < 0.001, **** *P* < 0.0001. *Created in BioRender. Schneider, N.* (*2026*) https://BioRender.com/lxc76kq

Interestingly, comparison of ASO_6, which was fully complementary to the region containing the target adenosine, with ASO_6C, which contained a C-to-A mismatch opposite the target nucleotide, revealed an approximately four-fold decrease in correctly spliced transcripts in uninduced HeLa cells (Supplementary Figure 4). This finding suggests that this mismatch, facilitating ADAR-based SDRE, may already partially alleviate splice disruption. We therefore concluded that this region of the original ex/ex gRNA can remain within the optimized gRNA design, while other disruptive regions of the gRNA should be modified.

We next designed three optimized gRNAs to mitigate splice-disruptive elements identified by the ASO walk, incorporating modifications in length and sequence (Figure 7C). These included a truncated 53-nt gRNA lacking the first three 5′ nucleotides and last four 3′ nucleotides (v50.3, 5’−25-C-27), a gRNA containing a G-to-A mismatch opposite the penultimate nucleotide of exon 11 (v50.4, 5’−29-C-2-G-27), and a third gRNA incorporating a G-to-U mismatch within the central region of the identified splice-disruptive motif (v50.5, 5’−16-G-12-C-30). Capillary electrophoresis analysis showed a significant improvement in splicing for two of the gRNAs with v50.3 gRNA and v50.5 gRNA showing similar most improved splicing overall (Figure 7D). Sanger sequencing analysis of the WT-spliced transcripts showed 33.1%, 51.0%, and 48.8% editing of the target adenosine for gRNAs v50.3, v50.4, and v50.5 respectively (Figure 7E). Overall, v50.5 gRNA achieved the most favorable combined outcome, increasing the proportion of edited WT-length transcripts from 21% to 30% (1.5-fold) out of all final transcripts (Figure 7F). Together, these results indicate that rational gRNA engineering can simultaneously mitigate splice interference while preserving substantial levels of on-target RNA editing.

## DISCUSSION

In this study, we investigated how gRNA design and transcript context influence both SDRE and pre-mRNA splicing, while evaluating the potential of ADAR-mediated editing as a strategy for splice modulation. This work was motivated by a gap in the field: although considerable effort has been devoted to optimizing ADAR-recruiting gRNAs to maximize on-target editing and minimize off-target editing activity (6–16), comparatively little attention has been paid to their effects on splicing or to the therapeutic correction of splice-altering mutations. Our findings both show that the majority of G>A P/LP variants are splice-related or in close proximity to a splice boundary and identify key design principles that should be considered to maximize the therapeutic potential of ADAR-recruiting gRNAs while minimizing unintended effects on splicing.

Although endogenous ADAR1 and 2 editing has been extensively documented in intronic sequences and in regions that become part of mature mRNA (61–64) to our knowledge no other study directly measured SDRE levels occurring on pre-mRNA vs mRNA separately. A previous study compared the editing efficiencies of pre-mRNA- and mRNA-targeting gRNAs using a symmetric linear 111-nt guide design (6) where the 5’ sequence of the gRNA was identical before the orphan cytosine, and subsequent 15 nucleotides downstream were identical as well before encountering the exon boundary. This might allow one gRNA to bind and edit both transcript types, since the regions immediately flanking the orphan cytosine remained identical between the corresponding pre-mRNA and mRNA. In another study, an exon-targeting gRNA was used for SDRE and indirectly measured editing of mRNA as the difference between pre-mRNA and subsequent mRNA editing levels (12). By contrast, we designed a system in which the target adenosine was positioned at the splice junction, enabling transcript-specific targeting of pre-mRNA or mature mRNA. To our knowledge, this is the first demonstration that pre-mRNA and mature mRNA can be selectively targeted by SDRE using transcript-specific gRNAs with minimal shared target recognition, enabling direct measurement of editing within each transcript type. We observed that pre-mRNA and mRNA can be differentially edited under these conditions, indicating that transcript context can influence SDRE efficiency. No consistent trend was observed favoring either pre-mRNA or mRNA across ADAR2-overexpressing conditions, suggesting that editing efficiency is largely sequence-dependent. Additionally, co-delivery of both gRNA types did not reduce editing levels or result in additive editing effects, consistent with competitive binding to the exon harboring the adenosine target. It is possible that half of the gRNA would bind to the exonic target, blocking the intended gRNA from binding and editing. Together, these findings highlight an important consideration for future SDRE studies. Because many proof-of-concept editing experiments are initially performed using minimal reporter constructs expressing the cDNA sequence, they may overlook transcript-context effects that become apparent in splice-competent systems. Incorporating minigene, midigene, or endogenous gene models earlier in the development pipeline may allow for more effective gRNA design with the target transcript-type in mind.

ADAR has previously been shown to affect splicing in an editing-independent manner, whereby binding of the ADAR protein to target RNA may impede access of the spliceosome (27–31). This characteristic may represent an important and underappreciated risk of "off-target" effects of SDRE, namely ADAR-mediated splice disruption. Consistent with this possibility, we observed a strong correlation between editing levels and splice interference for terminal exonic targets. Regardless of the gRNA target transcript type, higher editing levels were associated with lower levels of correctly spliced transcripts, suggesting that ADAR recruitment and/or prolonged occupancy of the target RNA may interfere with spliceosome assembly or function. Our findings also emphasize the importance of accurately quantifying RNA editing in the context of potential splice perturbation. Primers used to assess editing should not rely solely on amplification of the edited exon, as exon skipping may occur and confound measurements of editing efficiency while masking splice-disruptive effects of the gRNA.

We further observed evidence for the complex spatial and temporal relationship between SDRE and splicing when targeting exonic splice mutations in *CDH23* and *BBS1*. Editing levels were consistently higher in WT-length transcripts than in their intron-retained counterparts in both ADAR2-overexpressing and uninduced cells. However, up to 20% editing was still detected in intron-retaining transcripts, indicating that RNA editing can correct a splice mutation even in transcripts that ultimately undergo aberrant splicing. This suggests that ADAR-mediated editing and spliceosome activity may compete spatially and/or temporally. Nevertheless, for *CDH23* and *BBS1*, ADAR2 overexpression resulted in higher editing levels and reduced intron retention, demonstrating that ADAR-mediated SDRE can be harnessed to modulate splicing despite the dynamic interplay between RNA editing and spliceosome activity.

At least 16% of pathogenic variants causing inherited retinal diseases affect splicing (65) and we show here that ∼39% of all P/LP G>A mutations are splice-related, yet these types of variants remain largely unexplored as targets for SDRE. We therefore sought to evaluate the therapeutic potential of ADAR-mediated editing across several classes of splice-affecting variants. To our knowledge, only a limited number of studies have investigated this concept, including targeted editing of a terminal exonic splice mutation causing ornithine transcarbamylase deficiency and SDRE-induced exon skipping as a therapeutic strategy for Duchenne muscular dystrophy (25, 26). In the present study, we evaluated SDRE of a deep-intronic pseudoexon variant, a near-canonical (+5) donor splice-site variant, and exonic splice-affecting variants. For the deep-intronic pseudoexon, we found that simulated editing of the splice acceptor, rather than the mutant nucleotide itself, was sufficient to abolish pseudoexon inclusion, illustrating that neighboring splice-regulatory nucleotides may represent effective therapeutic targets. Consistent with the findings of Guo et al. (26), where gRNA introduction and ADAR1 expression in an ADAR-deficient cell line further enhanced exon skipping, pseudoexon skipping for this variant was greatest following gRNA delivery in ADAR1-overexpressing cells. In contrast, the +5 splice donor-region variant proved considerably more challenging to rescue. Although optimization of gRNA design successfully minimized splice interference, efficient restoration of normal splicing was not achieved, suggesting that intronic variants located immediately adjacent to the splice donor may present particular challenges for ADAR-mediated correction. This may partly explain the relative scarcity of splice-site-adjacent intronic SDRE studies reported to date. Finally, for both exonic splice-affecting variants, increased ADAR-mediated editing was accompanied by improved splicing, demonstrating that SDRE has the potential to not only correct pathogenic nucleotides but also to restore normal splicing.

Finally, our study demonstrates that additional design principles should be considered when optimizing gRNAs to minimize splice interference. By avoiding predicted ESE motifs and modifying gRNA symmetry, we achieved meaningful editing levels together with splice rescue for exonic splice mutations. We further established a proof-of-concept workflow for gRNA optimization using an ASO walk to identify otherwise difficult-to-detect splice-sensitive regions. Incorporating these findings into gRNA design through guide truncation and strategically placed mismatches reduced splice disruption, with the most effective optimized gRNA increasing the proportion of correctly spliced, edited transcripts by 1.5-fold. Together, these findings demonstrate that rational gRNA design can improve the therapeutic potential of SDRE by balancing efficient editing with preservation of normal splicing.

This study has several limitations. Our experiments were performed primarily using plasmid-based splice reporters in HeLa cells with exogenous ADAR overexpression and therefore do not fully recapitulate endogenous chromatin context, transcriptional kinetics, or nonsense-mediated decay. Future studies on patient-derived cells should evaluate these design principles at endogenous loci and in disease-relevant cell types to determine how transcript context influences SDRE under physiological conditions.

Our optimization strategy further suggests that splice-regulatory elements should become an integral component of SDRE guide design. While considerable effort has been devoted to minimizing bystander and off-target editing, comparable high-throughput approaches for identifying and avoiding splice-sensitive regions are currently lacking. Future design algorithms could incorporate predicted ESEs, together with other cis-regulatory elements including exonic splicing silencers (ESSs), intronic splicing enhancers (ISEs), and intronic splicing silencers (ISSs), to minimize splice disruption while preserving editing efficiency. Such approaches may become increasingly important as the field continues to adopt longer gRNAs (11–13), which have a greater probability of overlapping functional splice-regulatory motifs. By assessing the effects of SDRE on transcript specificity, splice variants, and coding variants, we show in this study that ADAR-mediated SDRE and splicing are deeply intertwined and that future SDRE design should not only increase on-target editing and reduce bystander editing, but preserve normal RNA processing as well.

## ACKNOWLEDGMENTS

The authors would like to thank Prof. Frans Cremers and Prof. Rob Collin for the generous gift of *ABCA4* midigene plasmids and Prof. Muayyad Al-Ubaidi for the 661W cell line. Figures were created using BioRender.com.

## AUTHOR CONTRIBUTIONS

Nina Schneider: Conceptualization, Methodology, Validation, Formal analysis, Investigation, Resources, Data curation, Writing – original draft, Writing – review & editing, Visualization, Project administration. Nave Zehoray: Data curation, Writing – original draft, Visualization. Ricky Steinberg: Methodology, Writing – review & editing. Eyal Banin: Resources, Funding acquisition, Writing – review & editing. Yvan Arsenijevic: Funding acquisition, Writing – review & editing. Shay Ben Aroya: Resources, Methodology, Writing – review & editing, Funding acquisition. Erez Y. Levanon: Methodology, Data curation, Writing – review & editing, Funding acquisition. Dror Sharon: Conceptualization, Methodology, Resources, Writing – original draft, Writing – review & editing, Supervision, Project administration, Funding acquisition

## SUPPLEMENTARY DATA STATEMENT

Supplementary Data are available online.

## ETHICS STATEMENT

This study did not involve human participants, human tissues, or animals requiring ethical approval. The study used established cell lines and publicly available genetic variant data.

## DATA AVAILABILITY

All processed data supporting the findings of this study are included in the article and its Supplementary Data. The source code used in this study (Figure 1 and Supplementary Table 1) is available on GitHub (https://github.com/znavezz/clinvar-ga-splicing) and is permanently archived in Zenodo at 10.5281/zenodo.21692997. Raw sequencing data, Sanger chromatograms, capillary electrophoresis data, and ddPCR output files are available from the corresponding author upon reasonable request.

