## Supplementary Information for "Transcript-Specific Site-Directed RNA Editing Reveals Principles for Splice-aware guide RNA Design"

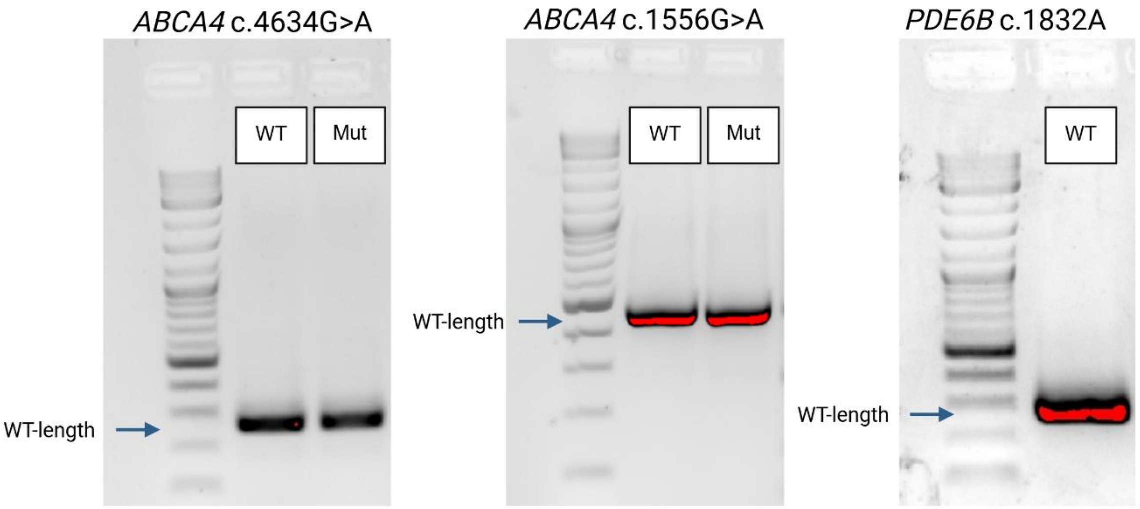


**Supplementary Figure 1**

Confirmation of correctly spliced wild-type-length products generated from splice constructs

PCR products demonstrating the expected correctly spliced wild-type (WT)-length transcripts generated from the *ABCA4* c.4634 WT, *ABCA4* c.4634G>A mutant (Mut), *ABCA4* c.1556 WT, *ABCA4* c.1556G>A mutant (Mut), and *PDE6B* c.1832 WT splice constructs.

A B

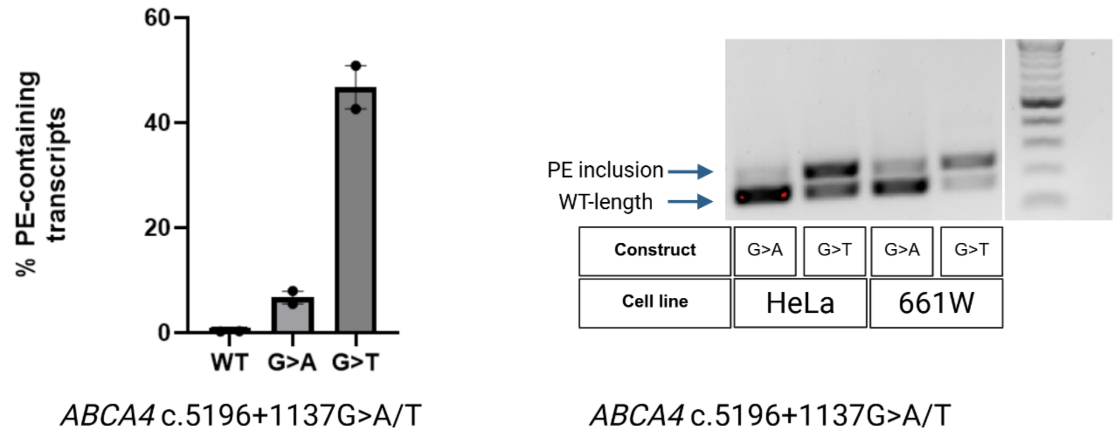


**Supplementary Figure 2**

The *ABCA4* c.5196+1137G>T variant results in greater pseudoexon inclusion than c.5196+1137G>A in HeLa and 661W cells

(A) ddPCR analysis of percent of pseudoexon-containing transcripts for wildtype (WT), *ABCA4* c.5196+1137G>A (G>A), and *ABCA4* c.5196+1137G>T (G>T) midigene plasmids (n=2 for each). (B) Agarose gel analysis of PCR products generated from *ABCA4* c.5196+1137G>T and c.5196+1137G>A splice constructs transfected into HeLa and 661W cells, demonstrating greater pseudoexon inclusion for the c.5196+1137G>T variant in both cell lines. PE, pseudoexon.

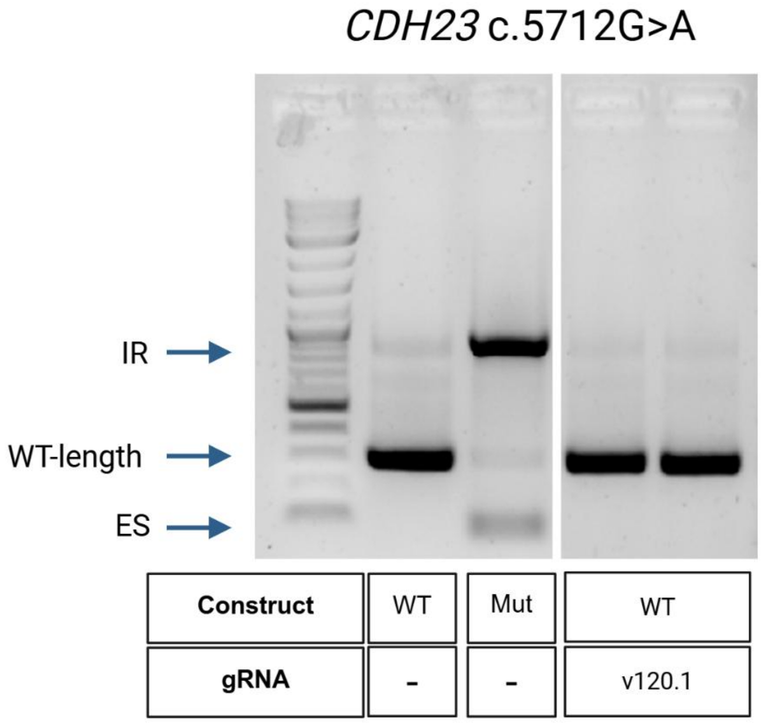


**Supplementary Figure 3**

gRNA v120.1 does not disrupt wild-type splicing

PCR analysis of splice products generated from the *CDH23* c.5712G wild-type (WT) splice construct with or without gRNA v120.1, alongside the *CDH23* c.5712G>A mutant (Mut) splice construct. IR, intron retention; ES, exon skipping.

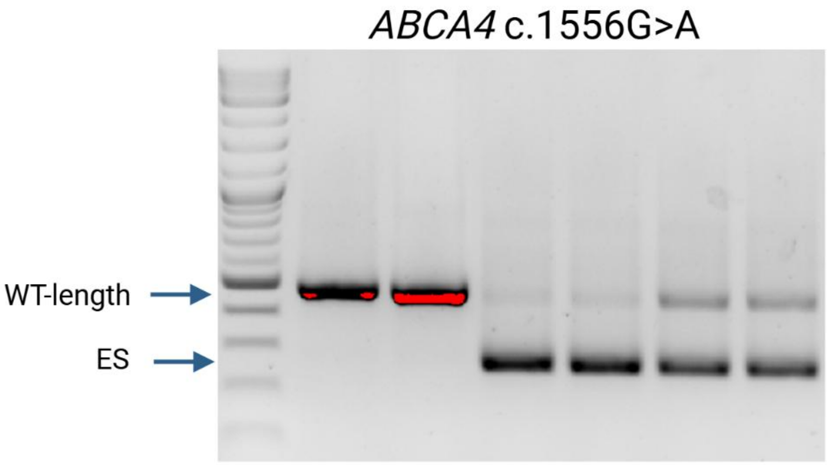


| Construct | WT | Mut | Mut | Mut |
|---|---|---|---|---|
| **AON** | - | - | AON6 | AON6C |
| **%WT-length transcripts** | 100% | 100% | 6.4% | 24.6% |

**Supplementary Figure 4**

Introduction of a C-to-A mismatch reduces AON6-induced missplicing of *ABCA4* c.1556G>A

PCR analysis of splice products generated from the ABCA4 c.1556G>A mutant (Mut) splice construct treated with AON6, AON6C, or left untreated, alongside the *ABCA4* c.1556G wild-type (WT) splice construct. ES, exon skipping. %WT-length transcripts calculated from capillary electrophoresis.

**Supplementary Table 1. Source values for G>A variant classification**

| Section | Metric | n | Out_of | Percent | Range |
|---|---|---|---|---|---|
| cohort | P/LP SNVs (exonic + intronic within MANE) | 188621 | | | |
| cohort | G>A variants analysed (Figure 1C and 1D) | 40552 | | | |
| cohort | G>A variants classified (Figure 1B) | 40437 | | | |
| cohort | excluded: intergenic / outside MANE | 468 | | | |
| cohort | excluded from Figure 1B: exonic G>A with no SpliceAI score | 115 | | | |
| cohort | variants with no SpliceAI score - any substitution | 985 | | | |
| distance | G>A within 0 nt of a junction | 1513 | 40552 | 3.70% | |
| distance | G>A within 1 nt of a junction | 15833 | 40552 | 39.00% | |
| distance | G>A within 30 nt of a junction | 25672 | 40552 | 63.30% | |
| distance | G>A within 50 nt of a junction | 30771 | 40552 | 75.90% | |
| distance | G>A within 100 nt of a junction | 36242 | 40552 | 89.40% | |
| distance | G>A within 500 nt of a junction | 39188 | 40552 | 96.60% | |
| splice classes | Donor site | 9491 | 40437 | 23.50% | |
| splice classes | Acceptor site | 5239 | 40437 | 13.00% | |
| splice classes | Intronic (splice region) | 160 | 40437 | 0.40% | |
| splice classes | Deep intronic | 170 | 40437 | 0.40% | |
| splice classes | Exonic (splicing suspected) | 524 | 40437 | 1.30% | 1.3%-1.6% |
| splice classes | Exonic | 24853 | 40437 | 61.50% | |
| splice classes | TOTAL splice-associated (every class except Exonic) | 15584 | 40437 | 38.50% | 38.5%-38.7% |
| deep intronic | Deep intronic - more than 30 nt into an intron - any substitution | 947 | 188621 | 0.50% | |
| deep intronic | Of those- G>A | 170 | 947 | 18.00% | |

Each row gives a count (n), its denominator (out_of) and the resulting percentage. Note the two denominators: 40,552 G>A variants for the distance rows and 40,437 for the splice classes, because 115 exonic G>A variants have no SpliceAI score. The range column bounds the values that depend on a SpliceAI prediction; an empty range means the value is exact.

### Supplementary Table 2. List of gRNA antisense sequences.

| gRNA | Symmetricity (5’-3’) | Variant | Synthesis scale | Purification | Sequence: r(RNA base), m (2'O-methyl, *(phosphothiorate bond) |
|---|---|---|---|---|---|
| **v21.1** | 29-C-30 | *ABCA4* c.4634G>A | 100 nmol | HPLC | mA*mU*mA*rArArArArGrCrArUrArArArArGrGrArUrUrUrCrUrUrCrUrUrArCrCrUrGrCrUrUrCrUrUrArUrArArGrArGrCrArGrGrArUrArCrGrUrUrU*mU*mU*mA |
| **v22.2** | 29-C-30 | *ABCA4* c.4634G>A | 100 nmol | HPLC | mG*mU*mU*rCrArUrUrGrArCrCrCrArGrArArUrUrUrGrCrUrCrUrUrUrArArGrCrUrGrCrUrUrCrUrUrArUrArArGrArGrCrArGrGrArUrArCrGrUrUrU*mU*mU*mA |
| **v50.1** | 29-C-30 | *ABCA4* c.1556G>A | 100 nmol | HPLC | mU*mG*mU*rArGrCrUrUrUrCrArArArCrUrUrArUrCrCrArGrGrArCrCrArArGrCrArCrUrGrCrArGrArGrArGrUrCrArCrArArArGrUrUrGrArGrArGrA*mG*mU*mG |
| **v50.2** | 29-C-30 | *ABCA4* c.1556G>A | 100 nmol | HPLC | mU*mG*mU*rArGrCrUrUrUrCrArArArCrUrUrArUrCrCrArGrGrArCrCrArArGrCrArCrUrCrCrArGrGrUrArUrUrGrArUrUrGrArCrCrArGrGrCrGrGrA*mG*mG*mG |
| **v70.1** | 29-C-30 | *PDE6B* c.1832A | 100 nmol | HPLC | mA*mA*mU*rUrCrArCrArUrGrCrCrCrGrCrCrCrUrGrArGrGrUrGrCrCrUrArCrCrUrCrArUrCrUrGrGrUrArCrArGrGrUrUrGrUrUrGrGrUrGrCrCrGrC*mG*mG*mU |
| **v70.2** | 29-C-30 | *PDE6B* c.1832A | 100 nmol | HPLC | mC*mG*mU*rGrGrArGrCrUrUrArGrCrCrArArGrGrGrGrUrUrCrUrGrGrGrArCrCrUrCrArUrCrUrGrGrUrArCrArGrGrUrUrGrUrUrGrGrUrGrCrCrGrC*mG*mG*mU |
| **v40.1** | 28-C-30 | *ABCA4*: c.5196+1137G>T | 2 nmol | Standard desalting | mU*mC*mC*rUrCrCrCrCrGrUrGrCrArUrUrUrArUrGrArGrUrGrUrUrUrCrCrCrArGrUrGrUrArGrArCrArGrArArGrArGrArGrArGrGrArArArUrArA*mU*mU*mU |
| **v27.1** | 29-C-30 | *ABCA4* c.5714+5G>A | 100 nmol | HPLC | mC*mU*mG*rCrCrCrArCrUrGrGrCrCrCrArGrGrGrUrGrUrGrGrCrArUrGrGrArCrGrUrArCrCrArUrUrGrGrGrArGrArGrGrArArGrArArGrUrGrGrCrG*mC*mU*mG |
| **v27.2** | 48-C-10 | *ABCA4* c.5714+5G>A | 2 nmol | Standard desalting | mC*mA*mG*rUrUrCrUrGrGrArUrGrCrCrCrUrGrArGrCrUrGrCrCrCrArCrUrGrGrCrCrCrArGrGrGrUrGrUrGrGrCrArUrGrGrArCrGrUrArCrCrArU*mU*mG*mG 3’ |
| **v27.3** | 28-C-10 | *ABCA4* c.5714+5G>A | 2 nmol | Standard desalting | mU*mG*mC*rCrCrArCrUrGrGrCrCrCrArGrGrGrUrGrUrGrGrCrArUrGrGrArCrGrUrArCrCrArU*mU*mG*mG 3’ |
| **v27.4** | 11-C-10 | *ABCA4* c.5714+5G>A | 100 nmol | Standard desalting | mU*mG*mU*rGrGrCrArUrGrGrArCrGrUrArCrCrArU*mU*mG*mG |

| | | | | | |
|---|---|---|---|---|---|
| **v27.5** | 37-C-9-CCUC-9 | *ABCA4* c.5714+5G>A | 100 nmol | HPLC | mG*mC*mC*rCrUrGrArGrCrUrGrCrCrCrArCrUrGrGrCrCrCrArGrGrGrUrGrUrGrGrCrArUrGrGrACrGrUrArCCrArUrUrGrCrCrUrCrArGrGrArArG*mA*mA*mG |
| **v120.1** | 40-C-19 | *CDH23* c.5712G>A | 100 nmol | HPLC | mG*mA*mA*rArCrCrCrUrGrGrGrArArGrCrUrCrCrCrUrGrGrGrGrUrCrArGrUrCrUrArGrGrCrCrCrUrArCrCrGrUrGrGrCrArUrUrGrArUrGrArArGrA*mA*mG*mG |
| **v130.1** | 41-C-18 | *BBS1* c.479G>A | 100 nmole | Standard desalting | mG*mA*mA*rGrGrArGrUrGrArGrGrGrGrGrGrUrArUrGrArArGrGrGrGrArArGrGrCrArGrCrCrUrCrUrCrArCrCrGrGrArUrGrCrUrCrUrCrCrArGrCrA*mU*mC*mU |
| **v50.3** | 25-C-27 | *ABCA4* c.1556G>A | 100 nmol | HPLC | mG*mC*mU*rUrUrCrArArArCrUrUrArUrCrCrArGrGrArCrCrArArGrCrArCrUrCrCrArGrGrUrArUrUrGrArUrUrGrArCrCrArGrGrC*mG*mG*mA |
| **v50.4** | 29-C-2-G-27 | *ABCA4* c.1556G>A | 100 nmol | HPLC | mU*mG*mU*rArGrCrUrUrUrCrArArArCrUrUrArUrCrCrArGrGrArCrCrArArGrCrArCrGrCrCrArGrGrUrArUrUrGrArUrUrGrArCrCrArGrGrCrGrGrA*mG*mG*mG |
| **v50.5** | 16-G-12-C-30 | *ABCA4* c.1556G>A | 100 nmol | HPLC | mU*mG*mU*rArGrCrUrUrUrCrArArArCrUrUrGrUrCrCrArGrGrArCrCrArArGrCrArCrUrCrCrArGrGrUrArUrUrGrArUrUrGrArCrCrArGrGrCrGrGrA*mG*mG*mG |

### Supplementary Table 3. List of gBlock sequences.

| | Gene | Variant | Sequence Type | Sequence 5'-3' | Length (bases) | Restriction enzymes | RNA scheme |
|---|---|---|---|---|---|---|---|
| **Minigene splice constructs** | *ABCA4* | c.1556G>A | WT | cagcaagagGGGCCCCTCGAGaatggctgaagaacaagaccaaagatcctat<br>ggaatttctaagcagagcagtgactgtatttcttcttcccaagGATACCCTGGGGAA<br>CCCAACAGTAAAAGACTTTTTGAATAGGCAGCTTGGTGAAGAA<br>GGTATTACTGCTGAAGCCATCCTAAACTTCCTCTACAAGGGC<br>CCTCGGGAAAGCCAGGCTGACGACATGGCCAACTTCGACTG<br>GAGGGACATATTTAACATCACTGATCGCACCCTCCGCCTGGT<br>CAATCAATACCTGGAGgtaaggggctgcaagccccacagtgggccccttgaag<br>atagccccatgagtggggccagagctcccttagcaagtcaagtggtcttgaatttaagctttc<br>attttccccactgaagaaacaagaatccctacatcccctgtacagttctcattctctaacagct<br>tatccatacttaaaacctctggagttaagcaaacattaagtttataggcctccttgcatttgacc<br>atttcctgggacagcagcccttatcctgtgactttctgtgtgtagagttgagtctttgcagttggtc<br>ctcctcacactctctcaactttgtgactctctgcagTGCTTGGTCCTGGATAAGTT<br>TGAAAGCTACAATGATGAAACTCAGCTCACCCAACGTGCCCT<br>CTCTCTACTGGAGGAAAACATGTTCTGGGCCGGAGTGGTATT<br>CCCTGACATGTATCCCTGGACCAGCTCTCTACCACCCCACGT<br>GAAGTATAAGATCCGAATGGACATAGACGTGGTGGAGAAAAC<br>CAATAAGATTAAAGACAGgtgatgtttcaggaagggctcgctgcatttctccaaa<br>gtcagtgggaaattacatttggtagagagaaagggattgagactggactcataaGGAT<br>CCACTAGtcaccttgtc | 943 | XhoI and BamHI_HF | partial intron 10-Exon 11-Intron 11 with shortened center-Exon 12- partial intron 12 |
| | *ABCA4* | c.1556G>A | Mutant | cagcaagagGGGCCCCTCGAGaatggctgaagaacaagaccaaagatcctat<br>ggaatttctaagcagagcagtgactgtatttcttcttcccaagGATACCCTGGGGAA<br>CCCAACAGTAAAAGACTTTTTGAATAGGCAGCTTGGTGAAGAA<br>GGTATTACTGCTGAAGCCATCCTAAACTTCCTCTACAAGGGC<br>CCTCGGGAAAGCCAGGCTGACGACATGGCCAACTTCGACTG<br>GAGGGACATATTTAACATCACTGATCGCACCCTCCGCCTGGT<br>CAATCAATACCTGGAGgtaaggggctgcaagccccacagtgggccccttgaag<br>atagccccatgagtggggccagagctcccttagcaagtcaagtggtcttgaatttaagctttc<br>attttccccactgaagaaacaagaatccctacatcccctgtacagttctcattctctaacagct<br>tatccatacttaaaacctctggagttaagcaaacattaagtttataggcctccttgcatttgacc<br>atttcctgggacagcagcccttatcctgtgactttctgtgtgtagagttgagtctttgcagttggtc<br>ctcctcacactctctcaactttgtgactctctgcagTACTTGGTCCTGGATAAGTT<br>TGAAAGCTACAATGATGAAACTCAGCTCACCCAACGTGCCCT<br>CTCTCTACTGGAGGAAAACATGTTCTGGGCCGGAGTGGTATT<br>CCCTGACATGTATCCCTGGACCAGCTCTCTACCACCCCACGT<br>GAAGTATAAGATCCGAATGGACATAGACGTGGTGGAGAAAAC<br>CAATAAGATTAAAGACAGgtgatgtttcaggaagggctcgctgcatttctccaaa<br>gtcagtgggaaattacatttggtagagagaaagggattgagactggactcataaGGAT<br>CCACTAGtcaccttgtc | 943 | XhoI and BamHI_HF | partial intron 10-Exon 11-Intron 11 with shortened center-Exon 12- partial intron 13 |

| | | | | | | | |
|---|---|---|---|---|---|---|---|
| | *CDH23* | c.5712G>A | WT | cagcaagagGGGCCCCTCGAGgcccctcctctgcctctgcctttcctgcttccctcat<br>catcctcttttcatcttcttgcctttttaatgttctttttcttctttcttcctctccccttttctctttcttctccat<br>gaccaactgcaccctccctccctccatgccgcccttggcgaacctcctcctcggttgccatg<br>cacaacatctgtcgctcttcctcccctccctcctccttctctgactggcccagATGCTGGT<br>GGGGATCCGGGTGCTGGACATCAACGACAACGACCCTGTGC<br>TGCTGAACCTGCCCATGAACATCACCATCAGCGAGAACAGCC<br>CTGTCTCCAGCTTTGTCGCCCATGTCCTGGCCAGTGACGCTG<br>ACAGTGGCTGCAATGCACGCCTCACCTTCAACATCACTGCGG<br>GCAACCGCGAGCGGGCCTTCTTCATCAATGCCACGgtagggccta<br>gactgaccccagggagcttcccagggtttccagtgaaaaagaggaccctgccctagagg<br>cttttctgctgccaccatctgggctccaccgggctcccgttcctgcactgggatgaggacacc<br>tctgtgggaagcatggaatcttggattcaccagtgaaccatctctggagtccacctagagcc<br>agccctgaagcttgtcatgtgcagagatggggccagagcctgtgggaccacttggtttaatc<br>aagacgcaccaaagttctcacttagcctcttgaggtgtctgtagGGATCCACTAGtca<br>ccttgtc | 740 | ApaI and SpeI | part intron 41-Exon 42-part intron 42 |
| | *CDH23* | c.5712G>A | Mutant | cagcaagagGGGCCCCTCGAGgcccctcctctgcctctgcctttcctgcttccctcat<br>catcctcttttcatcttcttgcctttttaatgttctttttcttctttcttcctctccccttttctctttcttctccat<br>gaccaactgcaccctccctccctccatgccgcccttggcgaacctcctcctcggttgccatg<br>cacaacatctgtcgctcttcctcccctccctcctccttctctgactggcccagATGCTGGT<br>GGGGATCCGGGTGCTGGACATCAACGACAACGACCCTGTGC<br>TGCTGAACCTGCCCATGAACATCACCATCAGCGAGAACAGCC<br>CTGTCTCCAGCTTTGTCGCCCATGTCCTGGCCAGTGACGCTG<br>ACAGTGGCTGCAATGCACGCCTCACCTTCAACATCACTGCGG<br>GCAACCGCGAGCGGGCCTTCTTCATCAATGCCACAgtagggccta<br>gactgaccccagggagcttcccagggtttccagtgaaaaagaggaccctgccctagagg<br>cttttctgctgccaccatctgggctccaccgggctcccgttcctgcactgggatgaggacacc<br>tctgtgggaagcatggaatcttggattcaccagtgaaccatctctggagtccacctagagcc<br>agccctgaagcttgtcatgtgcagagatggggccagagcctgtgggaccacttggtttaatc<br>aagacgcaccaaagttctcacttagcctcttgaggtgtctgtagGGATCCACTAGtca<br>ccttgtc | 740 | ApaI and SpeI | part intron 41-Exon 42-part intron 42 |
| | *BBS1* | c.479G>A | WT | cagcaagagGGGCCCCTCGAGatggccatgagactggattcagataaaggacg<br>tgagcttggtgctcagtgcagaggtaggctggcagggaggcagagaccaagaggtaccc<br>ctagaagtgtggagctgtctgggggtgtagacattgggtttcctgcctggcttgagctcacag<br>gctctcttcccacattgtcacagGACCGAATCGACCCCTTAACCCTGAAG<br>GAGATGCTGGAGAGCATCCGgtgagaggctgccttccccttcatacccccctc<br>actccttcatcccatctgagccccagggccccattcttccattcggctgccatgctggcccttt<br>ccttgcagGGAGACGGCAGAGGAGCCTTTGTCCATCCAGTCACT<br>CAGgtaaggaccctgtggagggccagggtttgggaggctccagggagaggaagcag<br>ccgccagaatgtgccagaatgatggaggagggcagcgaggccggggcctgcagatgc<br>cagttctctgtgtgtctagGTTTCTGCAGCTGGAGCTAAGTGAAATGGA<br>GGCATTTGTAAACCAACACAAGTCCAACTCCATCAAGCGGCA<br>Ggtaatacccccttctctttttatttccctgtaaaaaaatttacatttttttaaaagaccacttaac<br>atcagtggtaaattctataagcaaacccaaccagtatctctttgctatatcccttccagccGG<br>ATCCACTAGtcaccttgtc | 740 | XhoI and BamHI_HF | part intron 4- exon 5- intron 5- exon 6- intron 6-exon 7- part intron 7 |

| | | | | | | | |
|---|---|---|---|---|---|---|---|
| | *BBS1* | c.479G>A | Mutant | cagcaagagGGGCCCCTCGAGatggccatgagactggattcagataaaggacgtgagcttggtgctcagtgcagaggtaggctggcagggaggcagagaccaagaggtacccctagaagtgtggagctgtctgggggtgtagacattgggtttcctgcctggcttgagctcacaggctctcttcccacattgtcacagGACCGAATCGACCCCTTAACCCTGAAGGAGATGCTGGAGAGCATCCAgtgagaggctgccttccccttcatacccccctcactccttcatcccatctgagccccagggccccattcttccattcggctgccatgctggcccctttccttgcagGGAGACGGCAGAGGAGCCTTTGTCCATCCAGTCACTCAGgtaaggaccctgtggagggccagggtttgggaggctccagggagaggaagcagccgccagaatgtgccagaatgatggaggagggcagcgaggccggggcctgcagatgccagttctctgtgtgtctagGTTTCTGCAGCTGGAGCTAAGTGAAATGGAGGCATTTGTAAACCAACACAAGTCCAACTCCATCAAGCGGCAGgtaataccccccttctcttttttatttccctgtaaaaaaatttacatttttttaaaagaccacttaacatcagtggtaaattctataagcaaacccaaccagtatctctttgctatatcccttccagccGGATCCACTAGtcaccttgtc | 740 | XhoI and BamHI_HF | part intron 4- exon 5- intron 5- exon 6- intron 6-exon 7- part intron 7 |
| | *PDE6B* | c.1832A | WT | cagcaagagGGGCCCCTCGAGgtcggaggtccaacctccaacccgacgcctaggtcatcccaacccctcaccactccccaccctgctggagccaggaccggtgagcaaggtggccctgtctctacagACCGGCAAACTGAAGAGCTACTACACGGACCTGGAGGCCTTCGCCATGGTGACAGCCGGCCTGTGCCATGACATCGACCACCGCGGCACCAACAACCTGTACCAGATGAAgtaggcacctcagggcgggcatgtgaattagccctaaatcaactccacgcccttggcgtgaattaggcttcgcatagcaggctatgtagaaagtggagtccacggccaggccccgtactccagcactgtgggaggccaaggcgaggggattg…aaaaaaagctgtgtatggtggcgcacacctgtggtcccagctacttaggaggctgaggtgggaggatcacttgagcccaggaggtcaaggctgtattgagccatgattgcaccactgcactccagagtgggtgacagaggcagaccctgtctcaaaaaaaagaaagtggggcccatctgggggggctgcagagcgcagggtgggccaagggcaggtcccacgggcctcacctccaccacctgtgtaacagGTCCCAGAACCCCTTGGCTAAGCTCCACGGCTCCTCGATTTTGGAGCGGCACCACCTGGAGTTTGGGAAGTTCCTGCTCTCGGAGGAGgttggtatactcaccctcggtttctgctgtgggcgctggggacgcagcgtccgcaggacggcagggccgtatcctgcggagcagggttctgatgcagcgggtgagcactgggtgtgtgagcactgggggagggcggcagagaaggcggagggccgaggctgagggcaggtgcatcGGATCCACTAGtcaccttgtc | 924 | Apa I and SpeI | part intron 13- exon 14- shortened intron 14- exon 15- part exon 15 |
| **Minimal splicing constructs** | *ABCA4* | c.4634G>A | mutant ex/in | gaacggccacgagttcgagatcgagggcgagggccagcaagagGGTACCtaaaaacgtatcctgctcttataagaagcaagtaagaagaaatccttttatgctttttatGGATCCcaccttgtcgtgagcaagggcgaggagctgttcaccggggtggtgccc | 163 | KpnI and BamHI-HF | 60 bases exactly complementary to ex/in gRNA |

| | | | | | | | |
|---|---|---|---|---|---|---|---|
| | *ABCA4* | c.4634G>A | mutant ex/ex | gaacggccacgagttcgagatcgagggcgagggccagcaagagGGTACCAAtaaaaacgtatcctgctcttataagaagcaacttaaagagcaaattctgggtcaatgaacAGGATCCcaccttgtcgtgagcaagggcgaggagctgttcaccggggtggtgccc | 163 | KpnI and BamHI-HF | 60 bases exactly complementary to ex/ex gRNA |
| | *ABCA4* | c.5196+1137 G>T | mutant intron/ PE | gaacggccacgagttcgagatcgagggcgagggccagcaagagGGTACCCCTaaattatttcctctctcttctgtctacactaggaaacactcataaatgcacggggaggacGGATCCcaccttgtcgtgagcaagggcgaggagctgttcaccggggtggtgccc | 166 | KpnI and BamHI-HF | 60 bases exactly complementary to gRNA |
| **Editing simulation constructs** | *ABCA4* | c.5196+1137 G>T | WT | cagcaagagGGGCCCCaatttattctaggtgtcttgagtagtaaaatctcaacattttaagaccaacatgagcctccatttcatgtgatgataagatataccaactgatggagaccaacacaaatgaccttctcatccatggttttttaaaatgatggtgaatattggaattcctgaagatatgatttctatcttactcagcttagtaagcagctatcacttaacaatacaaaaccagagattatcagtagcaactaaattatttcctctctcttctgtctacacgaggaaacactcataaatgcacggggaggaggtcagaacctgaaagcctttctttggataagagcatcaactgcaggtaccacattggccctgtgatgctaatataaaaggagctaggcccaccggtaccgaaaagttacttagaaaagtgcggaggcttttaattttactttttttaaaagataagaaatagaatttacGGATCCcaccttgtc | 500 | ApaI and BamHI-HF | part intron 36-pseudoexon-part intron 36 |
| | *ABCA4* | c.5196+1137 G>T | c.5196 +1137 G>T | cagcaagagGGGCCCCaatttattctaggtgtcttgagtagtaaaatctcaacattttaagaccaacatgagcctccatttcatgtgatgataagatataccaactgatggagaccaacacaaatgaccttctcatccatggttttttaaaatgatggtgaatattggaattcctgaagatatgatttctatcttactcagcttagtaagcagctatcacttaacaatacaaaaccagagattatcagtagcaactaaattatttcctctctcttctgtctacactaggaaacactcataaatgcacggggaggaggtcagaacctgaaagcctttctttggataagagcatcaactgcaggtaccacattggccctgtgatgctaatataaaaggagctaggcccaccggtaccgaaaagttacttagaaaagtgcggaggcttttaattttactttttttaaaagataagaaatagaatttacGGATCCcaccttgtc | 500 | ApaI and BamHI-HF | part intron 36-pseudoexon-part intron 36 |
| | *ABCA4* | c.5196+1137 G>T | c.5196 +1137 _1138 GA>TG | cagcaagagGGGCCCCaatttattctaggtgtcttgagtagtaaaatctcaacattttaagaccaacatgagcctccatttcatgtgatgataagatataccaactgatggagaccaacacaaatgaccttctcatccatggttttttaaaatgatggtgaatattggaattcctgaagatatgatttctatcttactcagcttagtaagcagctatcacttaacaatacaaaaccagagattatcagtagcaactaaattatttcctctctcttctgtctacactgggaaacactcataaatgcacggggaggaggtcagaacctgaaagcctttctttggataagagcatcaactgcaggtaccacattggccctgtgatgctaatataaaaggagctaggcccaccggtaccgaaaagttacttagaaaagtgcggaggcttttaattttactttttttaaaagataagaaatagaatttacGGATCCcaccttgtc | 500 | ApaI and BamHI-HF | part intron 36-pseudoexon-part intron 36 |

**Supplementary Table 4. Antisense oligonucleotide (ASO) sequences used to identify splice-critical regions surrounding ABCA4 c.1556G>A**

| **ASO name** | **Sequence (5'-3', all 2'MOE bases)** |
|---|---|
| **ASO1** | GATTGACCAGGCGGAGGG |
| **ASO2** | ATTGATTGACCAGGCGGA |
| **ASO3** | GGTATTGATTGACCAGGC |
| **ASO4** | CCAGGTATTGATTGACCA |
| **ASO5** | CTCCAGGTATTGATTGA |
| **ASO6** | AAACTTATCCAGGACCAAGTA |
| **ASO7** | TTCAAACTTATCCAGGAC |
| **ASO8** | GCTTTCAAACTTATCCAG |
| **ASO9** | TGTAGCTTTCAAACTTATC |
| **ASO6C** | AAACTTATCCAGGACCAAGCA |
| **SON** | UCAGACAUGUAAGUACUACUAGC |

### Supplementary Table 5. Summary of splice constructs and primers used for splicing and RNA editing analysis

| | | | | | Splice analysis | Primers for splice analysis | | Seq type | Primers for editing analysis | | Special sequencing comments |
|---|---|---|---|---|---|---|---|---|---|---|---|
| **Gene** | **c.** | **p.** | **construct** | **Plasmid backbone** | | **Forward** | **Reverse** | | **Forward** | **Reverse** | |
| *ABCA4* | c.4634G>A | p.Ser1545Asn | Minimal splicing construct | Addgene #86639 | N/A | N/A | N/A | NGS | 6873 | 6874 | |
| *ABCA4* | c.4634G>A | p.Ser1545Asn | Midigene splice construct | BAC (CH17-325O16) | Capillary electrophoresis | 6771 | 6772 | NGS | 6771 | 6772 | |
| *ABCA4* | c.1556G>A | p.Cys519Tyr | Minigene splice construct | pET01 | Capillary electrophoresis | ETPR04 | ETPR05 | Sanger | ETPR04 | ETPR05 | Gel excised and sequenced: ETPR04 for WT-length |
| *PDE6B* | c.1832A | N/A | Minigene splice construct | pET01 | Capillary electrophoresis | ETPR04 | ETPR05 | NGS | 6883 | 6884 | |
| *ABCA4* | c.5196+1137G>T | N/A | Minimal splicing construct | Addgene #86639 | N/A | N/A | N/A | NGS | 6873 | 6874 | |
| *ABCA4* | c.5196+1137G>T | N/A | Midigene splice construct | BAC (CH17-325O16) | ddPCR | ddPCR-F | ddPCR-R | N/A | N/A | N/A | |
| *ABCA4* | c.5196+1137G>T | N/A | Editing simulation construct | pET01 | Capillary electrophoresis | ETPR02 | ETPR03 | N/A | N/A | N/A | |
| *ABCA4* | c.5714+5G>A | N/A | Midigene splice construct | BAC (CH17-325O16) | Capillary electrophoresis | 6769 | 6770 | N/A | N/A | N/A | |
| *CDH23* | c.5712G>A | p.Thr1904= | Minigene splice construct | pET01 | Capillary electrophoresis | ETPR04 | ETPR05 | Sanger | ETPR04 | ETPR05 | Gel excised and sequenced: ETPR04 for WT-length, ETPR04 for Intron retention |
| *BBS1* | c.479G>A | p.Arg160Gln | Minigene splice construct | pET01 | N/A | ETPR04 | ETPR05 | N/A | N/A | N/A | |
| | | | | | Capillary electrophoresis | 6977 | ETPR05 | Sanger | 6977 | ETPR05 | Gel excised and sequenced with 6977 for WT-length bands, PCR purification and sequenced with 7035 for intron retention bands |

**Supplementary Table 6. List of PCR and primer sequences used for splicing and RNA editing analysis**

| Primer name | Primer sequence 5'-3' |
|---|---|
| 6769 | TGAGGAAGCTGCTCATTGTC |
| 6770 | TGTTCTCGTCCTGTGAGCAG |
| 6771 | TCGTCGGCAGCGTCAGATGTGTATAAGAGACAGcgcagcacggaaattctaca |
| 6771 | TCGTCGGCAGCGTCAGATGTGTATAAGAGACAGcgcagcacggaaattctaca |
| 6772 | GTCTCGTGGGCTCGGAGATGTGTATAAGAGACAGctcgtctccgtcttggacac |
| 6772 | GTCTCGTGGGCTCGGAGATGTGTATAAGAGACAGctcgtctccgtcttggacac |
| 6873 | TCGTCGGCAGCGTCAGATGTGTATAAGAGACAGggcatggacgagctgtacaa |
| 6873 | TCGTCGGCAGCGTCAGATGTGTATAAGAGACAGggcatggacgagctgtacaa |
| 6874 | GTCTCGTGGGCTCGGAGATGTGTATAAGAGACAGgtccagctcgaccaggatg |
| 6874 | GTCTCGTGGGCTCGGAGATGTGTATAAGAGACAGgtccagctcgaccaggatg |
| 6883 | TCGTCGGCAGCGTCAGATGTGTATAAGAGACAGCTGTGCCATGACATCGACCA |
| 6884 | GTCTCGTGGGCTCGGAGATGTGTATAAGAGACAGCACGATGCCGCGCTTCTG |
| 6977 | ACCGAATCGACCCCTTAACC |
| 6977 | ACCGAATCGACCCCTTAACC |
| 7035 | AATGGAAGAATGGGGCCCTG |
| ETPR02 | GATGGATCCGCTTCCTGCCCC |
| ETPR03 | CTCCCGGGCCACCTCCAGTGCC |
| ETPR04 | GGATTCTTCTACACACCC |
| ETPR05 | TCCACCCAGCTCCAGTTG |

**Supplementary Table 7. List of ddPCR primer sequences and probes used for ABCA4 c.5196+1137G>T midigene analysis**

| **Name** | **ssRNA type** | **Sequence (5'-3')** |
|---|---|---|
| ddPCR-F | Primer | AGCGGGTGAACAAATCCAAG |
| ddPCR-R | Primer | GAATGACCGCCCATCCATAC |
| Pseudoexon probe | Probe | /56-FAM/AATGCACGG/ZEN/GGAGGAGGTCAGAACC/3IABkFQ/ |
| Ex36_Ex37 probe | Probe | /5HEX/TCTGGGACA/ZEN/TCATGAATTATTCCGTGAG/3IABkFQ/ |
